# Actin-related protein M1 (ARPM1) required for acrosome biogenesis and sperm function in mice

**DOI:** 10.1101/2025.03.27.645694

**Authors:** Andjela Kovacevic, Eva Ordziniak, Naila Umer, Lena Arevalo, Leo D. Hinterlang, Sanaz Ziaeipour, Sara Suvilla, Gina E. Merges, Hubert Schorle

## Abstract

Actin-related proteins (Arps) are a superfamily of proteins which share sequence similarities with conventional actin and are involved in different cellular processes. Actin-related protein M1 (ARPM1) also known as actin-related protein T3 (ACTRT3) is a testis-enriched Arp which can be found in the perinuclear theca (PT) of murine round and elongating spermatids. ARPM1 forming a complex with Profilin 3 (PFN3) is lost in *Pfn3*-deficient sperm. We generated a mouse model deficient for *Arpm1* and demonstrate that *Arpm1*^-/-^ male mice are subfertile, with morphological aberrations of the acrosome. During spermiogenesis, defects become apparent in Cap phase of acrosome biogenesis when abnormal acrosomal granules are observed. *Arpm1*-deficiency causes deregulation of GM130 and TGN46 suggestive of defects in *cis-* and *trans-*Golgi trafficking required for acrosome development. Co-immunoprecipitation revealed that ARPM1 interacts with the PT-specific proteins ACTRT1, ACTRT2, ACTL7A and the sperm surface protein ZPBP, additionally to its already shown interaction with PFN3. We propose that ARPM1 acts as a structural component of the PT contributing to the cytoskeletal network connecting acrosome and nucleus. In addition, ARPM1 mediates the localization of ZPBP to enable fertilization and it tethers PFN3 to properly regulate Golgi-related acrosome development.

## Introduction

Spermiogenesis is a complex process of differentiation of male germ cells into mature and functional spermatozoa that takes place in seminiferous tubules of mammalian testes. During spermiogenesis, spermatids undergo DNA condensation and vast cellular remodeling including flagellum assembly, acrosome formation and eviction of cytoplasmatic residue. Due to the unique morphology of mature sperm cells, a complex network of cytoskeletal proteins is involved in forming of the spermatid and maintenance of the cellular structure in mature sperm. An important part of sperm cytoskeletal scaffolding is the perinuclear theca (PT), which is a rigid protein layer resistant to ionic detergents and high salt buffer extractions (Longo and Cook, 1991). The PT is located between the outer nuclear membrane and inner acrosomal membrane and develops along with acrosome formation during spermiogenesis (Oko and Sutovsky, 2009). The postacrosomal part of the PT is located at the caudal region of mature sperm where it forms a goblet like structure named calyx (Oko and Sutovsky, 2009). In PT protein lysates many sperm-specific cytoskeletal proteins can be found instead of conventional cytoskeletal proteins, reflecting the unique morphology and function of the sperm cells (Oko and Sutovsky, 2009). Extensive proteomic analyses of murine and bovine PT lysates revealed 500–800 different proteins, demonstrating the molecular complexity of this cytoskeletal structure. PT-specific proteins such as CCIN, CYLC1, CYLC2, actin-capping proteins, as well as members of actin-like and actin-related protein families co-localize and interact to establish a complex three-dimensional scaffold of the sperm head (Zhang et al., 2022a; Zhang et al., 2022b; Schneider and Kovacevic et al., 2023). However, molecular interactions and precise location of these proteins are just starting to be understood. In this study we elucidate the role of actin-related protein M1 (ARPM1) in the PT cytoskeletal complex.

Actin-related protein M1 (ARPM1) also known as Actin-related protein T3 (ACTRT3) belongs to a large family of Actin-related proteins (Arps) which share sequence similarities with conventional actin. Members of the Arp family are conserved from yeast to human (Goodson and Hawse, 2002) and have various roles such as actin polymerization, dynein motor function, as well as chromatin remodeling within the nucleus (Blessing et al., 2004). In addition, Arps are also found in the cytoplasm where they form complexes with actin-binding proteins and serve as cytoskeletal regulators. While most Arps are detected ubiquitously, several members of the family, such as ACTL7A, ACTL7B and ARPM1 are expressed exclusively in testis (Hara et al., 2008; Merges et al., 2023). Hara et al. showed that ARPM1 localizes in mouse spermatids and mature sperm and was detected in both cytoplasmatic and nuclear fraction of testis protein extracts. Interestingly, in late round spermatids and early elongating spermatids, ARPM1 localizes in the subacrosomal layer, while in testicular sperm it is detected in the postacrosomal calyx region of the PT (Hara et al., 2008).

While in spermatids and mature sperm ARPM1 forms a complex with PFN3, it coprecipitates with many other unidentified proteins (Hara et al., 2008). In this study, we generated an *Arpm1*-deficient mouse line using CRISPR/Cas9 mediated gene editing. Male *Arpm1*^-/-^ mice were subfertile with morphological defects of the acrosome, while the fertility of Arpm1⁺^/-^ male mice was unaffected, with no visible defects of the acrosome. First alterations become apparent in the Cap-phase of acrosome development in *Arpm1*^-/-^ spermatids, when irregular and vacuolized acrosomal caps are detected. Further, *cis*-and *trans*-Golgi networks required for proper acrosome biogenesis are affected as their markers, GM130 and TGN46 respectively, are deregulated. Using CoIP analyses, we identified ACTRT1, ACTRT2, ACTL7A and ZPBP as interaction partners of ARPM1 and confirmed the interaction with PFN3. The results suggest that ARPM1 contributes to the sperm cytoskeleton and acts as a molecular glue connecting the acrosome to PT and nuclear envelope. Further, we speculate that ARPM1 helps to connect and locate PFN3 to PT, in order to exert its role in orchestrating autophagic flux during spermiogenesis.

## Results

### *Arpm1* is conserved among species

To understand the relevance of *Arpm1* gene in the context of male fertility in humans and other species, we performed evolutionary analysis of selective constraints on *ARPM1* across rodents and primates. We revealed that the coding sequence is conserved in general, with rodents showing slightly higher conservation (Fig. S1, Supplementary Table S1). The evolutionary rate (ω) of the entire sequence for the entire tree studied was significantly lower than one, at ω=0.23 (Supplementary Table S1). A model that allows for the computation of the evolutionary rate for primates and rodents separately shows a small but significant difference between the clades ω, even though the coding sequence is significantly conserved in both (Rodent: ω= 0.21; Primate: ω= 0.29). Next, we estimated the ω for each codon site across the entire tree to examine the selective pressures across the coding region in greater detail. The model allowing for positive selection on codon sites explained the data significantly better than the null model according to likelihood ratio analysis (Table S1). 75% of codon sites were assigned to the conserved site class, 24% to the neutral/relaxed constraint site class and 1% to the positive selection site class. Only one codon site class was significantly positively selected (Fig. S1, Table S1). The distribution of evolutionary rates over the sequence shows a generally conserved constraint with several peaks exhibiting faster evolutionary rates. These faster evolving hot spots could encode for functional protein regions that have an adaptive advantage (Fig. S1). Overall, these findings show that *ARPM1* is a conserved constraint in general, indicating that it has an essential function. These results indicate that ARPM1 might have similar role across species.

In order to determine the expression pattern of *ARPM1* in mice and men, we first analyzed single cell RNA-sequencing data of adult mouse testis from Lukassen et al. As shown in Fig. S2 A, *Arpm1* is only weakly expressed in early stages from spermatocytes to round spermatids and Leydig cells, while spermatogonia and Sertoli cells lack *Arpm1* RNA. *Arpm1* is significantly upregulated in elongating spermatids (ES) and in compacted sperm (CS) (Fig. S2 A). Data extracted from *Human testis atlas* of young adult human testicular tissue indicates that human *ARPM1* is highly expressed in round and elongating spermatids (Fig. S2 B) (Guo et al., 2018). Taken together, the expression pattern of *ARPM1* in mouse and human are highly similar and the sequence is strongly conserved among species, suggesting that it has an important role during spermiogenesis of rodents and primates.

### Generation of *Arpm1*-deficient mouse lines

To elucidate the role of ARPM1 in sperm development, we generated *Arpm1*-deficient mouse lines using CRISPR/Cas9 mediated gene editing. Two guide RNAs targeting exon 1 and exon 2 were designed causing a 1.7 kb deletion resulting in a frameshift and a nonfunctional, highly truncated protein (Fig. 1 A, B). Two independent founder animals (Δ2295 and Δ2298) were generated and mated to C57Bl/6J WT mice to establish the lines. Sequencing analysis revealed that approximately 1.7 kb of the sequence was deleted, causing a deletion of large parts of Exon 1 and Exon 2 generating a fusion of remaining sequence and a frameshift (Fig. 3 A, B, C). The predicted truncated proteins would be 84 and 80 amino acids in size, as compared to 369 aa for the WT (Fig. 1 B). Genotyping revealed a band at 315 bp (Δ2295) and 319 bp (Δ2298) indicative for the deleted allele, while the WT allele produced a band 272 bp in size (Fig. 1 C). Polymerase chain reaction on cDNA revealed presence of truncated *Arpm1* transcripts in testes of *Arpm1*^-/-^ mice, while longer transcripts were detected for the WT (Fig. S2 C). As *Arpm1*-deficient animals from Δ2295 and Δ2298 did not show phenotypical differences, both lines were used for further analysis. A western blot using a polyclonal antibody detected no ARPM1 protein in testes extracts of *Arpm1*^-/-^ males, while a band corresponding to ARPM1 protein in WT and Arpm1⁺^/-^ was seen (Fig. 1D). Immunohistochemical staining showed ARPM1 detectable across different populations of germ cells in WT and Arpm1⁺^/-^ testicular sections, while in *Arpm1*^-/-^ testis tissue sections ARPM1 staining was absent (Fig. 1E). Finally, immunofluorescence staining of mature sperm revealed that ARPM1 localizes mainly in the posterior region of the perinuclear theca - calyx (Fig. 1 F). As expected, ARPM1 signal was absent from *Arpm1*^-/-^ epididymal sperm (Fig. 1 F). These results strongly suggest nonsense-mediated RNA decay of the truncated transcripts detected in *Arpm1*^-/-^ mice and indicate, that the gene editing generated a null allele for ARPM1.

**Figure 1:**
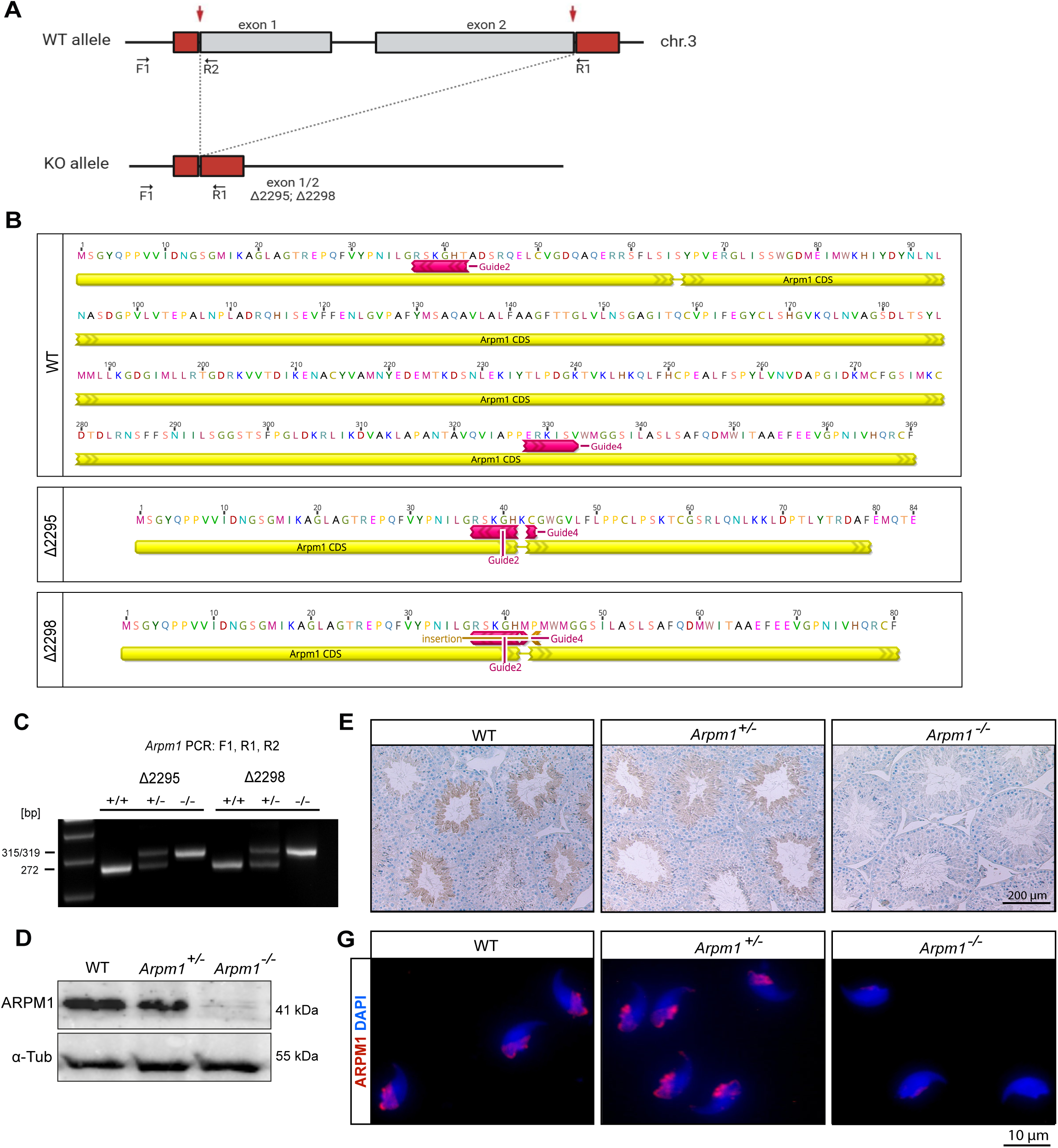
Generation of Arpm1-knockout mouse line **(A)** Schematic representation of the targeting strategy for CRISPR/Cas9-mediated generation of *Arpm1-*deficient alleles. Targeting sites of guide RNAs are depicted by red arrows. Genotyping primer binding sites are depicted by black arrows. **(B)** Schematic representation of the amino acid sequence of the truncated ARPM1 protein obtained in lines Δ2295 and Δ2298 compared to a WT amino acid sequence. Binding sites of guide RNA sites used are depicted in red. **(C)** Representative genotyping of WT, *Arpm1*^+/-^ and *Arpm1*^-/-^ mice by PCR. WT allele is detectable as a 272 bp PCR product, while *Arpm1*-knockout from lines Δ2295 and Δ2298 are observed as 315 bp and 319 bp PCR band, respectively. **(D)** Western Blot against ARPM1 on whole testis protein lysate of WT, *Arpm1* ^+/-^ and *Arpm1*^-/-^ mice. Band corresponding to ARPM1 is detected at 41 kDa. Alpha-tubulin (55 KDa) was used as load control. **(E)** Immunohistochemistry staining against ARPM1 on testicular tissue sections from WT, *Arpm1*^+/-^ and *Arpm1*^-/-^ male mice. Scale bar: 200 µm. Staining was performed on three animals of each genotype. **(F)** Immunofluorescent staining of WT, *Arpm1* ^+/-^ and *Arpm1*^-/-^ epididymal sperm cells against ARPM1 (red). Nuclei were counterstained with DAPI (blue). Scale bar: 10 µm. Staining was performed on three animals of each genotype.

### ARPM1 deficiency leads to male subfertility in mice and lower fertilization rate*in vitro*

Next, we performed fertility analysis to investigate the effect of *Arpm1*-deficiency on male fertility in mice. While pregnancy rates were not affected by the loss of ARPM1 (Fig. 2 A), *Arpm1*^-/-^ male mice were subfertile, with an average litter size of 1.5 compared to an average litter size of 7.5 in WT and Arpm1⁺^/-^ male mice mated with C57Bl/6J WT females (Fig. 2 B). Testis weight and overall morphology were not affected by *Arpm1* deficiency (Fig. 2 C, D). Histological analysis of Papanicolaou (PAS) stained testis tissue sections of all three genotypes revealed no alterations of seminiferous epithelium with all stages of spermatogenesis detectable (Fig. 2 E, Fig. S4 A). Further, in *Arpm1*^-/-^ male mice, a reduction of epididymal sperm was observed while, sperm count was not altered in Arpm1⁺^/-^ mice (Fig. 2 F). Eosin-Nigrosine staining of epididymal sperm, revealed that only 40% of sperm are viable in *Arpm1*^-/-^ mice compared to 80% in WT and Arpm1⁺^/-^ (Fig. 2 G, Fig. S4 B). The motility of the epididymal sperm activated in THY medium was not altered by *Arpm1*-deficiency (Fig. 2 H), suggesting that loss of ARPM1 does not affect the swimming ability of the sperm. Similarly, MITO-red staining of mitochondria revealed no alterations of the flagellum in Arpm1⁺^/-^ nor *Arpm1* ^-/-^ epididymal sperm, with regular distribution of mitochondria detectable in the midpiece (Fig. 2 I). Next, nuclear morphology of DAPI stained sperm heads was assessed. Sperm head shape was not altered in Arpm1⁺^/-^ when compared to WT, however, *Arpm1*^-/-^ sperm heads were slightly smaller in area (Fig. 2 J, Fig. S4 C). Finally, CMA3 staining showed no alterations in *Arpm1*-deficient sperm (Fig. 3 K, Fig. S4 D), suggesting that, despite its localization in the sperm nucleus, lack of ARPM1 does not affect the protamination and DNA compaction.

**Figure 2:**
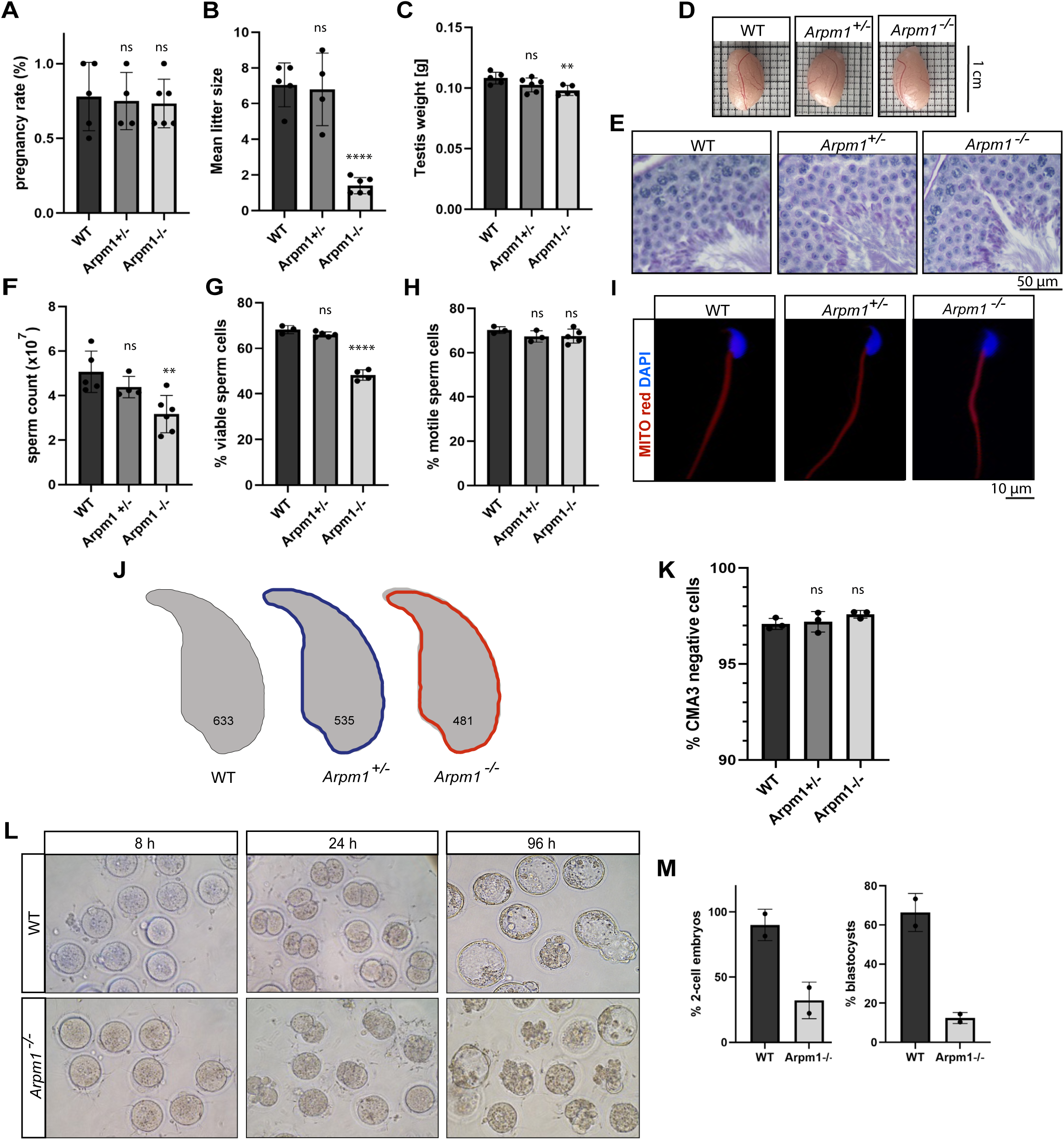
Arpm1-deficiency leads to male subfertility in mice. **(A)** Pregnancy rates (%) of adult WT (n=5), *Arpm1*^+/-^ (n=4) and *Arpm1*^-/-^ (n=6) male mice. Columns represent mean values ± standard deviation (s.d.). Black dots represent mean values obtained for each male animal included in the analysis. **(B)** Mean litter sizes of adult WT (n=5), *Arpm1*^+/-^ (n=4) and *Arpm1*^-/-^ (n=6) male mice. Columns represent mean values ± standard deviation (s.d.). Blac k dots represent mean values obtained for each male animal included in the analysis. **(C)** Average testis weight (g) of adult WT, *Arpm1*^+/-^ and *Arpm1*^-/-^ male mice. Columns represent mean values ± standard deviation (s.d.). Black dots represent mean values obtained for each male animal included in the analysis. **(D)** Comparable photographs of the testes of WT, *Arpm1*^+/-^ and *Arpm1*^-/-^ mice. Length of the box side: 1 cm **(E)** PAS staining of testicular tissue sections from WT, *Arpm1*^+/-^ and *Arpm1*^-/-^ mice. Scale bar: 50 µm. Staining was performed on three animals of each genotype. **(F)** Epididymal sperm count (×10^7^) of WT (n=5), *Arpm1*^+/-^ (n=4) and *Arpm1*^-/-^ (n=6). Columns represent mean values ± standard deviation (s.d.). Black dots represent mean values obtained for each male animal included in the analysis. **(G)** Quantification of viable sperm from WT (n=3), *Arpm1*^+/-^ (n=4) and *Arpm1*^-/-^ (n=4) male mice assessed with EN staining. Columns represent mean values ± standard deviation (s.d.). Black dots represent mean values obtained for each male animal included in the analysis. A minimum of 100 cells was counted for each sample. **(H)** Quantification of motile epididymal sperm from WT (n=3), *Arpm1*^+/-^ (n=3) and *Arpm1*^-/-^ (n=5) mice after activation in TYH medium. Columns represent mean values ± standard deviation (s.d.). Black dots represent mean values obtained for each male animal included in the analysis. **(I)** MITO red staining of epididymal sperm from WT, *Arpm1* ^+/-^ and *Arpm1*^-/-^ male mice showing intact flagella in all genotypes. Nuclei were counterstained with DAPI. Scale bar: 10 µm Staining was performed on three animals of each genotype. **(J)** Nuclear morphology analysis of *Arpm1*^+/-^ (blue outline) and *Arpm1* ^-/-^ (red outline) sperm heads compared to WT (gray). Number of cells analyzed for each genotype is shown. **(K)** Quantification of CMA3 negative sperm cells from WT, *Arpm1*^+/-^ and *Arpm1*^-/-^ male mice indicating normal DNA compaction. Black dots represent mean values obtained for each male animal included in the analysis. n=3. A minimum of 100 cells was counted for each sample. **(L)** An *in vitro f*ertilization assay on sperm from *Arpm1*^-/-^ and WT mice. Photographs of embryos at 8, 24 and 96 hours after fertilization are shown. **(M)** The percentage of two-cell embryos and percentage of blastocysts obtained during IVF experiment. Columns represent mean values ± standard deviation (s.d.). n=2 Statistical significance was calculated using a two-tailed, unpaired Student’s *t*-test. (\**P*<0.05, \*\**P*<0.005, \*\*\**P*<0.001, \*\*\*\**P*<0.0001).

**Figure 3:**
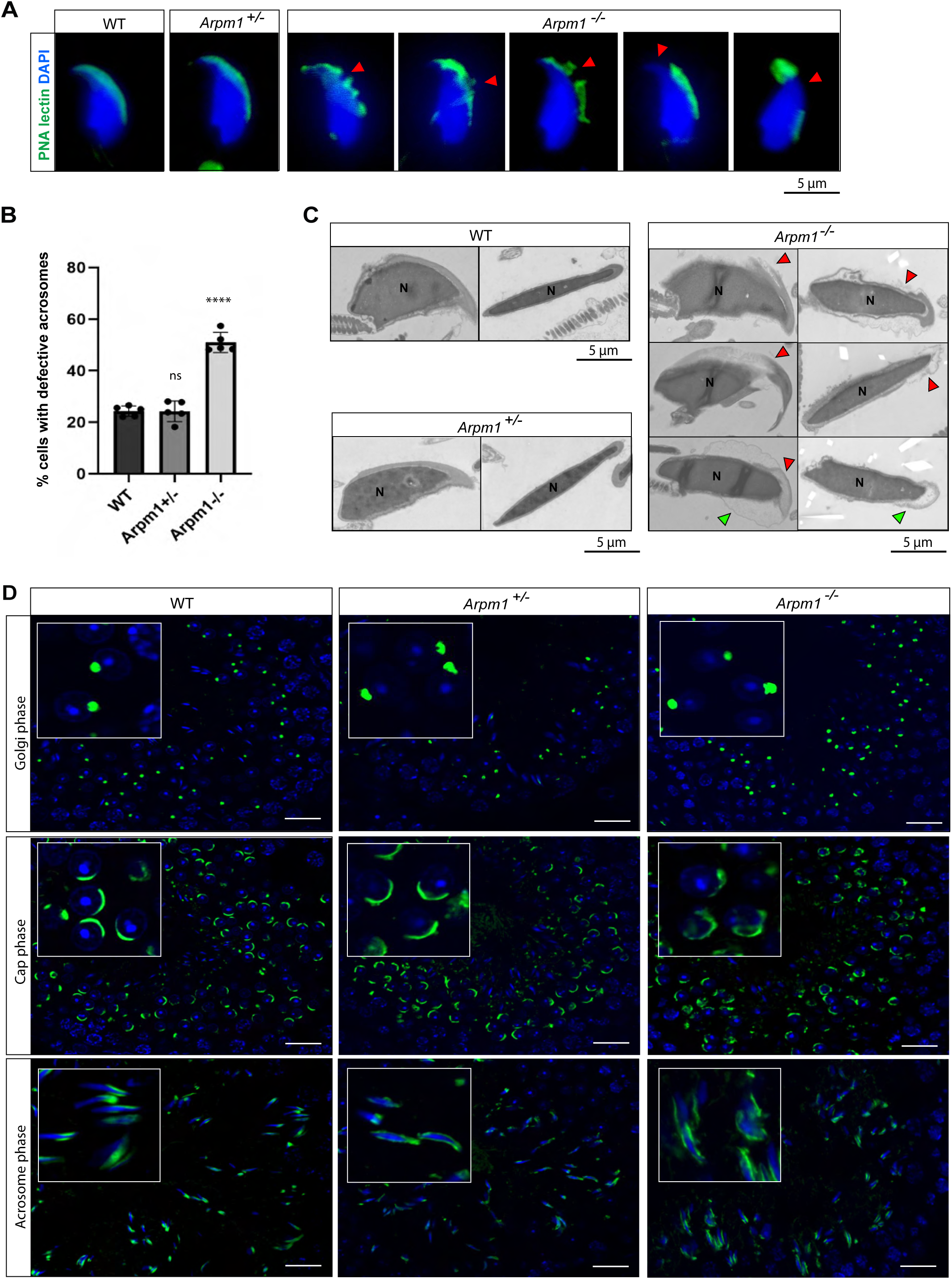
ARPM1 is required for acrosome integrity. **(A)** Immunofluorescence staining of epididymal sperm from WT, *Arpm1*^+/-^ and *Arpm1*^-/-^ male mice with Peanut agglutinin PNA-lectin (green). Nuclei were counterstained with DAPI (blue). Different acrosome defects in *Arpm1*^-/-^ sperm are shown; acrosome breakages are depicted with red arrowheads. Scale bar: 5 µm **(B)** Quantification of acrosome abnormalities observed in PNA-lectin-stained mature sperm of WT, *Arpm1*^+/-^ and *Arpm1*^-/-^ male mice. Columns represent mean values ± standard deviation (s.d.). Statistical significance was calculated using a two-tailed, unpaired Student‘s t-test. (\**P*<0.05, \*\**P*<0.005, \*\*\**P*<0.001, \*\*\*\**P*<0.0001). n=5. **(C)** Transmission electron microscopy (TEM) micrographs of WT, *Arpm1* ^+/-^ and *Arpm1*^-/-^ epididymal sperm. Defective acrosomes appear smaller on the apical portion of the sperm head and detached from the nucleus in *Arpm1*^-/-^ sperm (red arrowheads). Incorrectly evicted cytoplasm residues are depicted with green arrowheads. N: nucleus. Scale bar: 1 µm. **(D)** PNA-lectin immunofluorescence staining of the acrosome in testicular tissue sections of WT, *Arpm1*^+/-^ and *Arpm1*^-/-^ male mice (green). In Golgi phase of acrosome biogenesis at round spermatid stage acrosomal granule is visible by green dot-like signal at the surface of nuclei (upper panel). In Cap phases of WT and *Arpm1*^+/-^ round spermatids acrosomal granule is visible as a cap-like signal on one side of round spermatids (middle panel). In *Arpm1* ^-/-^ round spermatids, vacuolized acrosome granules are observed (middle panel, insert). The acrosomal phase is shown in the lower panel, with detached acrosomes visible in *Arpm1*^-/-^ elongating spermatids. Nuclei were counterstained with DAPI. Staining was performed on three animals from each genotype. Scale bar: 20 µm. Inserts show representative spermatids present in the main picture at higher magnification.

To investigate whether ARPM1 is required for fertilization and early embryo development we performed *in vitro* fertilization of WT oocytes using *Arpm1*^-/-^ and WT sperm. After 24 h of incubation in IVF medium, WT sperm had fertilized 90% of oocytes out of which 60% reached blastocyst stage (Fig. 2 L, M). Interestingly, *Arpm1*^-/-^ sperm displayed lower fertilization rate with only 35% of eggs reaching two-cell stage. Of note, blastocyst rate in relation to the two cell stage embryos remained comparable (65%) in both WT and *Arpm1-*deficient conditions. Sperm cells incubated in IVF medium showed normal swimming behavior and managed to find and bind to oocytes, however, they remained attached to the zona pellucida of the oocyte. This demonstrates that loss of *Arpm1* interferes with the ability of the sperm to penetrate the zona pellucida. The fact that fertilized oocytes progress to blastocyst stage at comparable speed and rates indicates, that ARPM1 is not required post fertilization. However, the observed fertilization deficiency suggests a defect in the acrosome reaction.

### ARPM1 deficient spermatids have defective acrosome development

Next, we analyzed acrosome morphology, biogenesis and function in detail. PNA-FITC staining of mature sperm showed that the acrosomes of *Arpm1*^-/-^sperm appeared vacuolized, shortened or not attached to the nuclear membrane (Fig. 3 A). Quantification of the PNA-FITC staining revealed that approximately 50% of *Arpm1* deficient sperm displayed acrosome abnormalities (Fig. 3 B). Staining against the acrosomal matrix marker, SP56 confirmed that 50% of *Arpm1*^-/-^ epididymal sperm had damaged acrosomes (Fig. S5 A, B). Transmission electron microscopy (TEM) of epididymal sperm showed that in the equatorial region of the *Arpm1*^-/-^ sperm head most acrosomes were partially detached from nuclear envelope (Fig. 3 C). On the apical pole of *Arpm1*^-/-^ sperm, acrosomes appear normal but drastically shortened or vacuolized, indicating that the acrosomes localize properly, however, the anchoring of the acrosome at the equatorial region appeared disturbed (Fig. 3 C, Fig. S5 C). As expected, acrosomes from *Arpm1*^+/-^ epididymal sperm were intact and comparable to the WT (Fig. 3 C, Fig. S5 C).

In order to determine when the observed defects originate, we performed PNA-FITC staining of testicular tissue to analyze acrosome development. In all three genotypes, in Golgi phase proacrosomal granules were clearly detectable on poles of round spermatids (Fig. 3 D). In WT and Arpm1⁺^/-^ testis, during Cap phase acrosomal granules form cap-like structures on the surface of round spermatids. However, in *Arpm1*^-/-^ spermatids, these cap structures do not form properly and appear disrupted, with the signal unevenly scattered among spermatids, resulting in vacuolization or detachment from the nuclear surface (Fig. 3 D; Fig. S5 D). During maturation phase, the acrosomal vesicles of WT and *Arpm1*^+/-^ elongating spermatids flatten over the nuclear surface on the apical side of the cell forming the arrow-like structures. However, in *Arpm1*^-/-^ testis sections the acrosomes appear thinner in the middle region of the cell and detached from the nucleus, similarly to what was observed in mature *Arpm1*^-/-^ sperm (Fig. 3 D; Fig. S5 E). Next, we performed TEM of testicular tissue to analyze in detail the ultrastructure during acrosome development. In round spermatids at step 4-5 regular acrosomal vesicles corresponding to Golgi phase were observed (Fig. S6 A). In Cap phase of *Arpm1*^-/-^ spermatids the Golgi apparatus appeared detached and displaced from the nuclear envelope, with large vesicles between the growing acrosome and the Golgi apparatus (Fig. 4 A). Since the vesicle trafficking between Golgi and the developing acrosome is essential for the proper acrosome development, this suggests that lack of ARPM1 eventually disrupts Golgi trafficking causing defects in the acrosome. Interestingly, round spermatids from Arpm1⁺^/-^ mice showed regular acrosomal granules with normal Golgi apparatus (Fig. S6 B). Of note, elongating spermatids of all three genotypes showed proper nuclear compaction and manchette formation, indicating that lack of ARPM1 does not interfere with DNA hypercondensation. However, in *Arpm1*^-/-^ spermatids, an increased gap between manchette and developing acrosome can be observed at the equatorial ring (Fig. 4B, Fig. S6 C).

**Figure 4:**
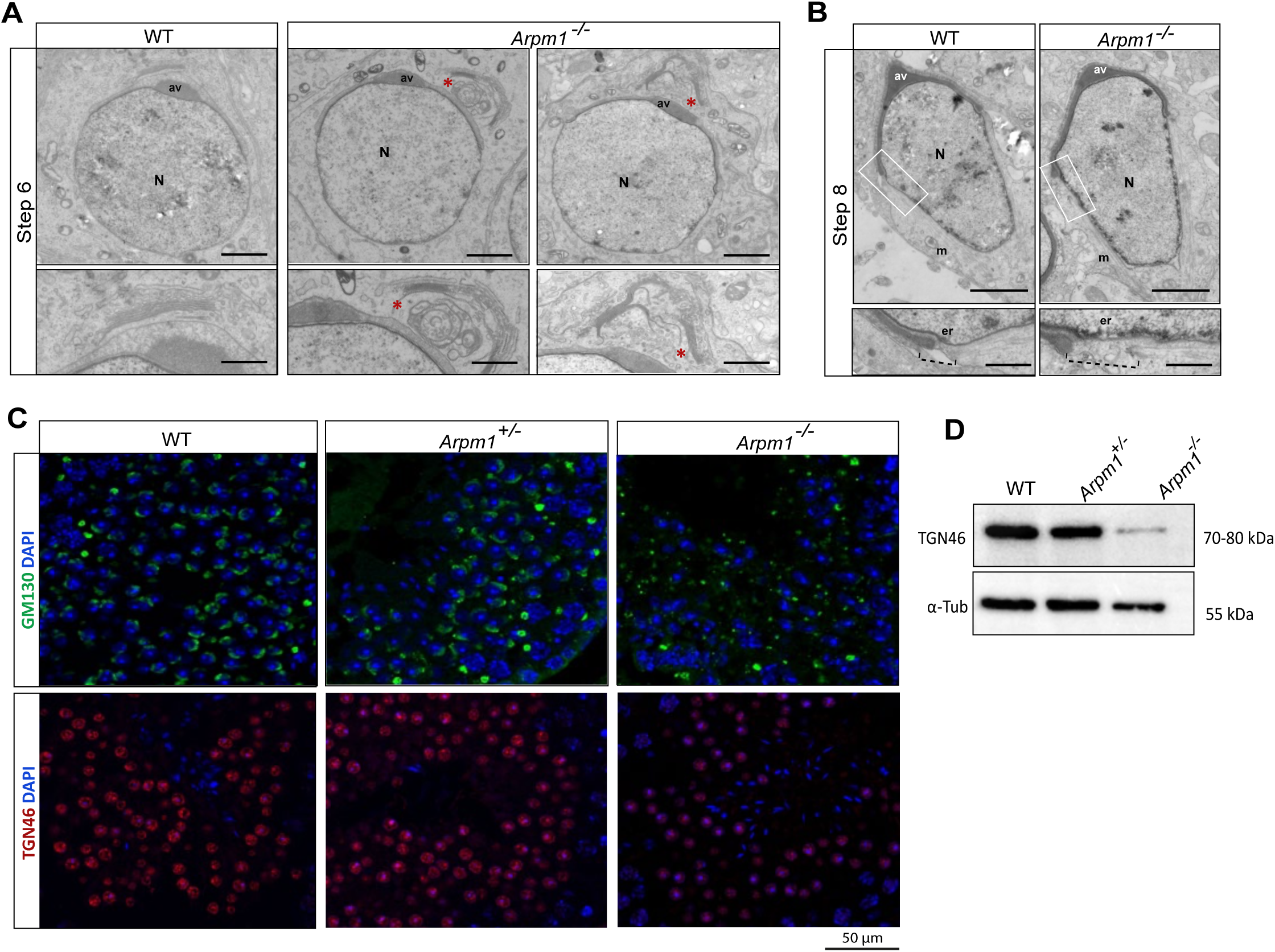
Arpm1 deficiency affects cis-and trans-Golgi networks. **(A)** Transmission electron microscopy (TEM) micrographs of WT and *Arpm1*^-/-^ round spermatids at step 6. In *Arpm1* ^-/-^ spermatids, Golgi apparatus is displaced and shows enlarged vacuoles depicted with asterisks. N: nucleus; av: acrosomal vesicle. Scale bar: 1 µm; insert scale bar: 5 µm **(B)** Transmission electron microscopy (TEM) micrographs of WT and *Arpm1*^-/-^ elongating spermatids at step 8. In *Arpm1*^-/-^ spermatids, enlarged space between acrosome and the manchette is observed at the equatorial ring (dashed line). N: nucleus; av: acrosomal vesicle; m= manchette; er: equatorial ring. Scale bar: 1 µm; insert scale bar: 5 µm **(C)** Immunofluorescence staining of testicular tissue sections from WT, *Arpm1*^+/-^ and *Arpm1*^-/-^ male mice against GM130 *cis*-Golgi marker (green, upper panel) and TGN46 *trans*-Golgi marker (red, lower panel). In WT and *Arpm1*^+/-^ round spermatids, GM130 signal forms a cap-like structure along the developing acrosome. In *Arpm1*^-/-^ round spermatids GM130 signal was scattered irregularly. In *Arpm1* ^-/-^ round spermatids TGN46 signal was less intense compared to WT and *Arpm1*^+/-^ round spermatids. Nuclei were counterstained with DAPI (blue). Scale bar: 50 µm. Staining was performed on three animals of each genotype. **(D)** Immunoblot against TGN46 on protein lysates from WT, *Arpm1*^+/-^ and *Arpm1*^-/-^ testes. The band is detected between 70-80 kDa. α-Tubulin was used as load control (55 kDa).

### *Arpm1*-deficient spermatids show defective Golgi trafficking

Since TEM analyses suggested that *Arpm1*-deficiency impinges on Golgi trafficking and/or vesicle tethering, we next analyzed *cis-* and *trans-*Golgi network markers. GM130 is required for vesicle tethering and maintenance of the *cis*-Golgi structural integrity and it presents as a cap-like staining pattern on round spermatids, corresponding to the growing acrosome as seen in WT and Arpm1⁺^/-^ testis (Tiwari et al., 2019). Interestingly, in *Arpm1*^-/-^ spermatids, the GM130 signal appeared irregularly dispersed, indicating that *Arpm1*-deficiency leads to aberrant *cis-*Golgi trafficking during acrosome biogenesis (Fig. 4 C, upper panel; Fig. S6 C). TGN46 is a *trans*-Golgi network marker required for the formation of exocytic vesicles and secretion from the *trans-*part of the Golgi network in round spermatids. In testicular tissue sections of WT and *Arpm1*⁺^/-^ mice, TGN46 is uniformly detected throughout round spermatids (Fig. 4 C, lower panel). However, in *Arpm1*^-/-^ testis tissues, the staining pattern appeared irregular with signal scattered around spermatids and with lower intensity (Fig. 4 C, lower panel; Fig. S6 D). Furthermore, western blot analyses revealed that protein levels of TGN46 were reduced in *Arpm1*^-/-^ testis compared to WT and Arpm1⁺^/-^ (Fig. 4 D). These results indicate that acrosome malformations in ARPM1-deficient mice are caused by defects in Golgi-derived proacrosomal vesicles. Of note, similar results were observed in *Pfn3*-deficient male mice (Umer et al., 2021), strongly suggesting that the PFN3-ARPM1 protein complex regulates Golgi-trafficking, which is required for acrosome biogenesis.

### ARPM1 interacts with other perinuclear theca proteins

As reported previously, ARPM1 interacts with PFN3 and co-precipitates with many other so far unidentified proteins (Hara et al., 2008). The PT consists of numerous cytoskeletal proteins, which closely interact to give rise to the complex 3D scaffold of the calyx. Based on its localization in the PT of spermatids and mature sperm, we hypothesized that ARPM1 might be a part of a protein complex that provides a cytoskeletal scaffold for the sperm head. We generated expression plasmids of various PT-specific proteins used in an *in vitro* system to study interactions between ARPM1 and other PT proteins. Combinations of expression plasmids containing the coding sequence of interest either HA-or Myc-tagged were transfected transiently in HEK293T cells. Immunofluorescent staining of transfected cells demonstrated the transfection efficiency (Fig. S7 A-G). Non-transfected HEK239T cells were stained with the same antibodies, and no signal was detected (Fig. S7 H).

First, co-immunoprecipitation (CoIP) confirmed the interaction between ARPM1 and PFN3. Furthermore, we revealed that ARPM1 interacts with ACTRT1 and ACTRT2, members of Arp family with testis-specific expression pattern. Interestingly, ARPM1 co-immunoprecipitates with ACTL7A which has been previously shown to interact with many other PT-proteins and is required for male fertility. Finally, ARPM1 co-immunoprecipitates with Zona pellucida binding protein (ZPBP), a sperm surface protein required for the binding to the zona pellucida of the oocyte. CoIP with ARPM1 and SPEM2 revealed no interaction (Fig. 5 A). SPEM2 is a testis-specific protein required for acrosome biogenesis and cytoplasm eviction (Li et al., 2024).

**Figure 5:**
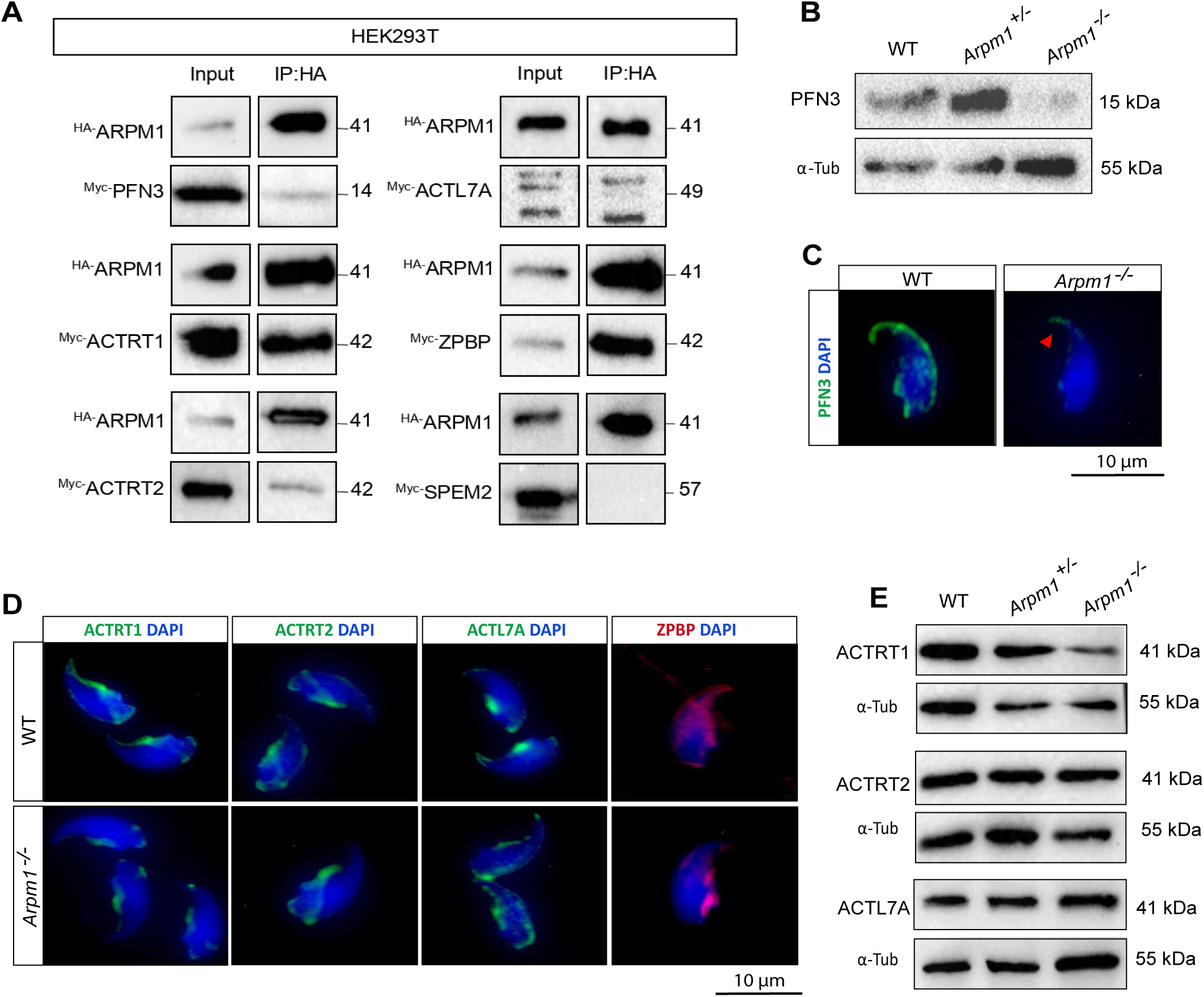
ARPM1 interacts with other PT proteins. **(A)** Co-IP assay of proteins expressed in HEK293T cells indicates an interaction between ARPM1 and PFN3, ACTRT1, ACTRT2, ACTL7A and ZPBP. The interaction between ARPM1 and SPEM2 was not detected. **(B)** Western blot against PT-specific proteins ACTRT1, ACTRT2 and ACTL7A on whole testis protein lysate of WT, *Arpm1*^+/-^ and *Arpm1*^-/-^ mice. Bands corresponding to ACTRT1, ACTRT2 and ACTL7A are detected around 40 kDa. α-tubulin (55 KDa) was used as load control. **(C)** Immunofluorescence staining against PFN3 in WT and *Arpm1*^-/-^ epididymal sperm. Red arrowhead depicts remaining PFN3 signal at the tip of the *Arpm1*^-/-^ sperm head. Nuclei were counterstained with DAPI. Scale bar: 10µm. **(D)** Immunofluorescence staining of PT-specific proteins that interact with ARPM1 in WT and *Arpm1*^-/-^ epididymal sperm. Localization of ACTRT1 and ACTRT2 is unaltered in *Arpm1*^-/-^ sperm. ACTL7A still localizes in the PT of *Arpm1*^-/-^ sperm, but with irregular distribution compared to WT. ZPBP accumulates in the calyx region of *Arpm1*^-/-^ epididymal sperm instead of acrosomal portion as seen in WT sperm. Nuclei were counterstained with DAPI. Scale bar: 10µm. Staining was performed on three animals of each genotype. **(E)** Westen Blot against PT-specific proteins ACTRT1, ACTRT2 and ACTL7A on whole testis protein lysate of WT, *Arpm1*^+/-^ and *Arpm1*^-/-^ mice. Bands corresponding to ACTRT1, ACTRT2 and ACTL7A are detected around 40 kDa. α-tubulin (55 KDa) was used as load control.

### ARPM1 is part of complex PT protein network

In order to further analyze whether lack of *Arpm1* effects its interaction partners in the PT we investigated their localization by immunohistochemical staining (IHC) and western blot. First, western blot analysis revealed that the level of PFN3 in *Arpm1*^-/-^ testicular protein extracts was dramatically reduced (Fig. 5 B, Fig. S8 B). This reduction of PFN3 was confirmed by IHC of testicular tissue sections from *Arpm1*^-/-^ mice (Fig. S8 A). In WT epididymal sperm, PFN3 signal can be detected throughout the PT surrounding the nucleus, while in samples from *Arpm1*^-/-^ mice, PFN3 signal is reduced and observed only at the tip of the sperm head (Fig. 5 C, red arrowhead). Interestingly, *Pfn3*-deficient mice displayed loss of ARPM1 in testicular lysates as well (Umer et al., 2021).

Next, we performed IHC against ACTRT1, which is located in the calyx of mature sperm in WT mice. In *Arpm1*^-/-^ sperm, ACTRT1 is detected in the calyx as well, however, the signal appeared weaker suggesting a lower protein abundance (Fig. 5 D). Similarly, reduced protein levels of ACTRT1 were detected on western blot of *Arpm1*^-/-^ testicular protein lysates compared to WT and *Arpm1*^+/-^ (Fig. 5 E, Fig. S8 C). Interestingly, *Arpm1*-deficiency did not affect neither the localization of ACTRT2 in the calyx of epididymal sperm (Fig. 5 D) nor its protein levels in testicular lysates (Fig. 5 E, Fig. S8 D). ACTL7A is located in the PT of WT sperm, with a typical accumulation in the region of equatorial ring. In *Arpm1*^-/-^ sperm, ACTL7A appears equally distributed along the PT, without the equatorial ring accumulation visible (Fig. 5 D). Interestingly, ACTL7A protein levels remained unaltered, suggesting that ARPM1 might be required for the proper localization of ACTL7A, but *Arpm1*-deficiency does not affect protein level (Fig. 5 D, Fig. S8 E). Finally, *Arpm1*-deficiency lead to a mislocalization of Zona pellucida binding protein (ZPBP) required for sperm to oocyte binding and fertilization (Lin et al., 2007). ZPBP localizes in acrosomal region of the sperm however in *Arpm1*^-/-^ sperm it accumulates in the posterior part of the sperm head (Fig. 5 D).

## Discussion

Here we demonstrate that *Arpm1* is highly conserved among species and expressed exclusively in the male germ line during spermiogenesis. *Arpm1*-deficient male mice are subfertile with defective acrosome development starting at the Cap phase, leading to irregular formed acrosomes in epididymal sperm. We demonstrate that ARPM1 interacts with PT proteins ACTL7A, ACTRT1, ACTRT2, PFN3, as well as the sperm surface protein ZPBP. These results indicate that in addition to being a structural protein within PT, ARPM1 helps to properly locate ZPBP in order to facilitate fertilization and it tethers PFN3 to the correct location which enables Golgi-mediated acrosome development.

We and others demonstrated the interaction between ARPM1 and PFN3 (Hara et al., 2008). Compared to the effects caused by loss of PFN3, the defects observed upon lack of ARPM1 are more confined. While *Pfn3*^-/-^ mice show defects in acrosome development, impaired sperm motility due to misshapen flagellae and cytoplasmic removal defects, loss of ARPM1 causes defects in acrosome development and shaping. Since we demonstrated that APRM1 interacts with PFN3 and PT proteins, we speculate that ARPM1 is required to localize PFN3 to the correct part within the developing acrosomal vesicles. Hence, additional defects observed in PFN3 deficient mice are effects independent of ARPM1.

Our results show that impaired *cis-* and *trans-*Golgi trafficking, as indicated by the disruption of GM130 and TGN46 respectively, contributed to alterations in acrosome biogenesis starting from round spermatids in *Arpm1*^-/-^ mice. Of note, impaired acrosomal granule formation and Golgi-trafficking during acrosome development were observed in *Pfn3*-deficient mice as well (Umer et al., 2021). However, acrosomal defects in *Pfn3*-deficient spermatids initiate at Golgi phase, which occurs earlier compared to the defects observed upon loss of ARPM1 detected during Cap phase. This could be explained by the slightly different RNA and protein expression patterns where PFN3 is detected at earlier stages of spermiogenesis compared to ARPM1 (Guo et al., 2018; Lukassen et al., 2018).

Furthermore, ARPM1 was lost in spermatids from *Pfn3*-deficient mice (Umer et al., 2021). In *Arpm1*-deficient testicular tissue and mature sperm, PFN3 protein levels were strongly reduced, suggesting that PFN3 and ARPM1 are not stable individually. Again, loss of PFN3 caused a more severe defect, that is lack of ARPM1, while deletion of ARPM1 diminished PFN3 levels. Thus, we speculate that PFN3 might also form complexes and be stabilized by ARPM1 related proteins such as ACTRT1 or ACTRT2.

Aside from PFN3 we demonstrated that ARPM1 interacts with PT proteins ACTRT1, ACTRT2 and ACTL7A. Of note, ACTRT1, ACTRT2 and ACTL7A are testis-enriched members of the family of Actin related proteins. ACTRT1 and ACTL7A are known to be part of a complex cytoskeletal network required for the proper development of the sperm and acrosome, as well as for the maintenance of the sperm morphology and function (Xin et al., 2020; Zhang et al., 2022b). Similar to our findings, male mice deficient for ACTRT1 are subfertile with defects of the acrosome and detachment from the nuclear envelope (Zhang et al., 2022b). Generation of ARPM1/ACTRT1 double deficient male mice could reveal, whether the proteins have redundant roles. Of note, *Actl7a*-deficient male mice are infertile with sperm showing severely altered head morphology, acrosome defects and detachment from the nuclear envelope (Xin et al., 2020; Zhou et al., 2022). ACTL7A has been found to interact with further PT-proteins ACTRT1 (Zhang et al., 2022b), ACTL9 (Dai et al., 2022), CCIN (Zhang et al., 2022a), CYLC1 (Jin et al., 2024) and SPACA1 (Chen et al., 2021) making it a central component of the complex protein network within the PT. Other than interaction with ACTL7A, ACTRT1 has been found to co-precipitate with ACTL9 and ACTRT2 (Zhang et al., 2022b) forming an ARP complex within the PT. Since ARPM1 interacts with ACTL7A and its interaction partner ACTRT1, we speculate that an actin-related complex acts as an integral part of the PT scaffold (Fig. 6).

**Figure 6:**
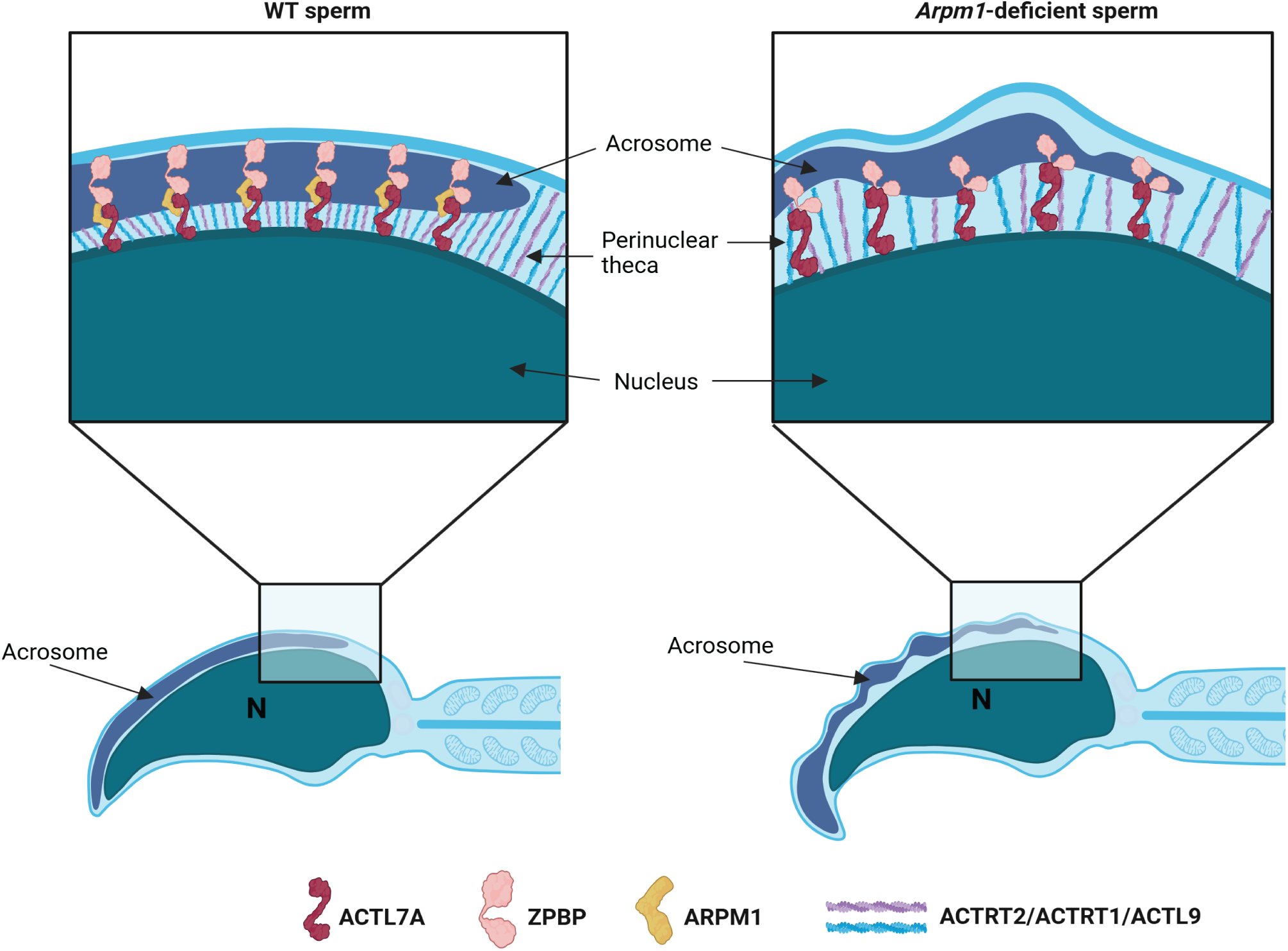
ARPM1 is a part of PT cytoskeletal scaffold. Schematic representation of PT network in WT and *Arpm1*^-/-^ epididymal sperm. Created in BioRender. Schorle, H. (2025) https://BioRender.com/153bjuu

Furthermore, ZPBP - a sperm-specific protein localized in the acrosome and required for binding and penetration of the zona pellucida - interacts with ARPM1 and is mislocalized in *Arpm1*-deficient sperm. This indicates that ARPM1 stabilizes ZPBP in its proper localization within the sperm head (Lin et al., 2007). We speculate that the mislocalization of ZPBP contributes to the subfertility of *Arpm1* ^-/-^ males.

Human patients carrying variants in *ACTL7A* or *ACTRT1* gene are infertile, with abnormal sperm head and acrosome morphology (Sha et al., 2021; Xin et al., 2020). Our evolutionary analysis shows that *ARPM1* is strongly conserved, suggesting that it might serve the same function in humans as well. Although human patients carrying variants in *ARPM1* gene are not identified yet, we speculate that it causes sub-or infertility in humans as well.

## Methods

### Animals

All animal experiments were conducted according to German law of animal protection and in agreement with the approval of the local institutional animal care committees (Landesamt für Natur, Umwelt und Verbraucherschutz, North Rhine-Westphalia, approval IDs: AZ84-02.04.2013.A429, AZ81-02.04.2018.A369). *Arpm1*-deficient mice were generated by CRISPR/Cas9-mediated gene-editing in zygotes of the hybrid strain B6D2F1. Guide sequences are depicted in Table 1. Two mouse lines were established and registered with Mouse Genome Informatics: B6-*Actrt3*^em1Hsc^ MGI:6718284 named line 2295 (NM_029690.3:c.125_995del) and line 2298 (NM_029690.3:c.125_993delinsTGCCTA). For all analyses sexually mature males at the age of 8-16 weeks, backcross generation ≥N3 were used.

**Table 1:**
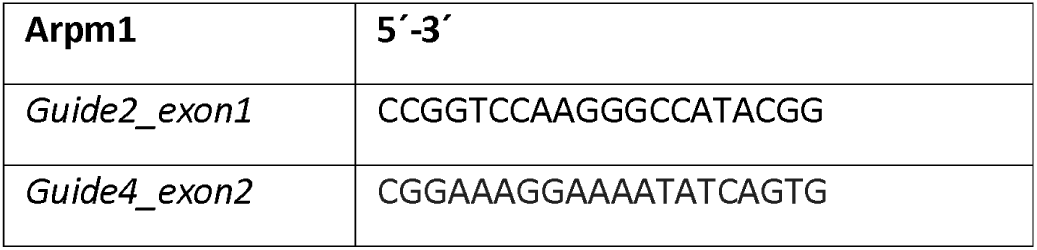
Guide sequences.

### Genomic DNA extraction, PCR genotyping and sequencing

Genomic DNA was extracted from biopsies using the HotShot method (Truett et al., 2000). PCR reactions were assembled according to the manufacturers protocol of the DreamTaq Green DNA Polymerase (Thermo Fisher, EP0712) using gene-specific primers listed in Table 3. PCR cycling conditions: 2 min 94°C; 35x (30 sec 94°C; 30 sec 66°C; 45 sec 72°C); 5 min 72°C. PCR products were separated on an agarose gel. PCR products or bands cut from agarose gels were, cleaned up and prepared for sequencing as described previously (Merges et al., 2022). Sequencing was performed with GATC/Eurofins and Genewiz (Azenta Life Sciences), using primers listed in table 2.

**Table 2:**
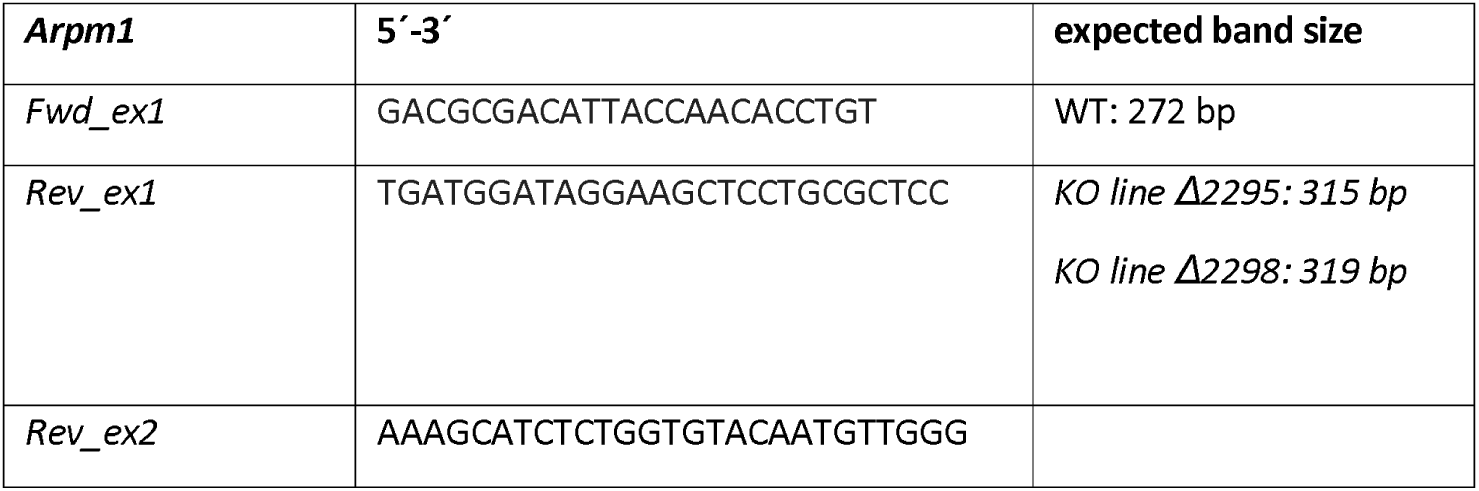
Primer sequences used in PCR genotyping and sequencing.

**Table 3:**
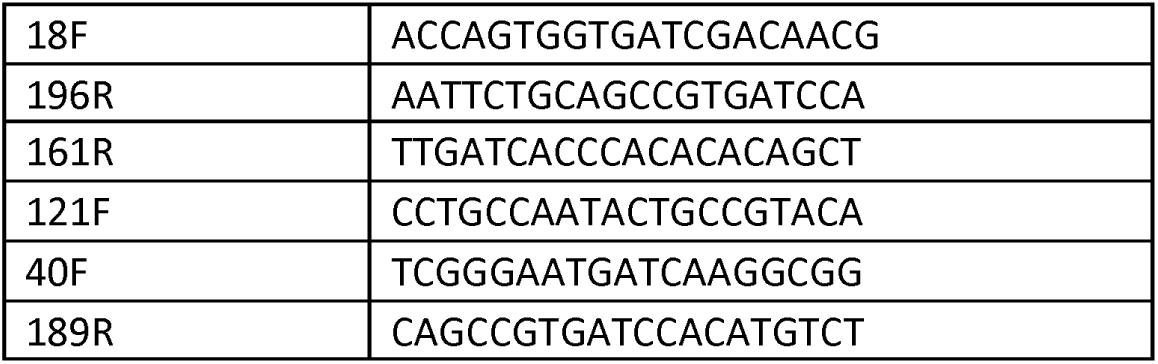
Primer sequences used in cDNA PCR.

### Fertility analysis and IVF

For fertility analysis male WT, Arpm1⁺^/-^ and *Arpm1*^-/-^ mice were mated 1:2/1:1 with C57Bl/6J females. Female mice were checked daily for the presence of a vaginal plug that indicates successful mating. Plug positive females were separated and pregnancies as well as litter size were recorded. For *in vitro* fertilization assay, C57Bl/6J WT females were super ovulated at 12 weeks of age. Experiment was performed twice, using WT and *Arpm1*^-/-^ male mice. Sperm extracted from both cauda epididymis were incubated in 200 µL of FertiUp and distributed among drops containing oocytes. The number of oocytes and embryos was monitored at 0h, 24h, 48h, 72h and 96h after fertilization.

### Sampling

Adult male mice were sacrificed at age of 8-16 weeks, and male reproductive organs were dissected. Both testes from each animal were weighted and fixed in Bouin or 4% PFA solution, or snap frozen, depending on further use. Cauda epididymides were collected in PBS preheated at 37°C and following several incisions of the cauda, sperm were retrieved by flush-out for 15-30 min. The extracted sperm samples were washed in PBS, centrifuged (4000 rpm, 5 min, 4°C) and snap frozen or fixed with methanol and acetic acid (3:1) solution, depending on the further use.

### Sperm concentration, viability and motility analysis

Epididymal sperm cells obtained by flush-out method were used for sperm counts using a Neubauer hemocytometer. Sperm concentration was analyzed for at least five animals for each genotype. Eosin-Nigrosin staining was used to determine sperm viability, as described previously (Schneider et al., 2020). Sperm motility was determined using epididymal sperm previously activated in THY medium. For all analyses at least 100 sperm per individual were analyzed and the percentage of motile/immotile, viable/inviable sperm was calculated.

### Sperm nuclear morphology analysis

To analyze sperm nuclear morphology, epididymal sperm cells from three animals of each genotype were fixed in methanol and acetic acid (3:1) and spread onto a glass slide. Next, sperm cells were stained with 4’,6-diamidino-2-phenylindole (DAPI) containing mounting medium (ROTImount FluorCare DAPI (Carl Roth GmbH, Karlsruhe, Germany; HP20.1)). The samples were imaged at ×100 magnification, using a Leica DM5500 B fluorescent microscope. At least 100 pictures were taken from each group and analyzed using Nuclear Morphology software according to the developer’s instructions (Skinner et al., 2019). The minimum detection area was 1000 pixels while the maximum detection area was 7000 pixels.

### RNA extraction and polymerase chain reaction on cDNA

Testicular RNA was extracted from two animals from each genotype using TRIzol (Life Technologies, Carlsbad, CA, USA; 15596018), according to manufactureŕs instructions. RNA concentrations and purity ratios were measured at NanoDrop ONE (Thermo Scientific). cDNA was transcribed using the Maxima H Minus Reverse Transcriptase Kit according to manufactureŕs instructions and oligo(dT) primers (Thermo Fisher Scientific). Approximately 100 ng cDNA was used in each PCR reaction. Primer used are listed in **Table 3**:

Different primer combinations were used to amplify potential transcripts: A:18F + 196R; B:18F + 161R; C: 121F + 196R; D:40F + 189R. Cycling conditions: **A** and **D**: 95°C, 60 sec; 30x (95°C, 30 sec; 60°C, 30 sec; 72°C, 30 sec); 5 min 72°C. **B** and **C**: 95°C, 60 sec; 30x (95°C, 30 sec; 60°C, 30 sec; 72°C, 45 sec); 5 min 72°C. A graphical representation of the targeted sequences as well as expected amplicon sizes are depicted in Fig. S2 C.

### Histology

Bouin or 4% PFA fixed mouse testis tissues were washed in 70% ethanol, paraffinized, embedded and sectioned at 3-5 µm using microtome. For histological analysis, the sections were deparaffinized with Xylene twice for 10 minutes and hydrated in alcohol row (5 minutes each). Tissue sections were then incubated with Periodic acid (0.5%) for 10 min, rinsed with H_2_0 and incubated for 20 min with Schiff reagent. After staining, the sections were dehydrated in alcohol row and mounted with Entellan (Sigma-Aldrich). Slides were imaged at 40x magnification under bright field using 5500 B microscope.

### Protein extraction

For protein extraction from testis tissues, the whole testis was homogenized in 1:10 RIPA buffer (1 ml per 100 mg of tissue), supplemented with Protease Inhibitors. Similarly, HEK293T cells were harvested in 500 µL DMEM and centrifuged at 1400 rpm, at 4°C for 4 minutes followed resuspension in RIPA buffer 1:10. After 15 minutes incubation on ice, samples were sonicated using Bioruptor (UCD-200TM-EX). Proteins for Co-immunoprecipitation experiments were isolated using M-PER™ Mammalian Protein Extraction Reagent (Thermo Scientific) supplemented with Protease inhibitors according to manufacturer’s instructions. Lysates were centrifuged at 14 000 g for 10 min and supernatant was collected. Samples were centrifuged for 30 minutes at 14 000 rpm, at 4°C. Protein concentrations were measured using Pierce™ BCA Protein Assay Kit (Thermo Fisher, #23225), according to manufactureŕs instructions for 96 well-plate approach. Absorption increases of purple reaction product corresponding to protein concentrations were measured using BioRad iMARK™ Plate Reader.

### Western Blot analysis

Protein extracts were separated on 10-12% SDS gel with 5% stacking gel. Trans Blot Turbo System (Bio-Rad) was used to transfer proteins to PVDF membranes. After washing with TBST 1X and staining with Coomassie blue, blots were blocked with 1% or 3% milk for 1 h at room temperature with gentle shaking. Alternatively, blocking was performed in 3% BSA for 1 h at room temperature. Primary antibodies were diluted in respective blocking solutions and incubated overnight at 4°C (antibody dilutions are listed in table 4). After washing in TBST, membranes were incubated for 1 h at room temperature with polyclonal goat anti-rabbit secondary antibody IgG/HRP (P044801-2; Agilent Technologies/Dako, Santa Clara, 527 CA, United States) or polyclonal rabbit anti-mouse secondary antibody IgG/HRP (P0260; Agilent Technologies/Dako) both diluted 1:2000 in blocking solution. Membranes were imaged using WESTARNOVA2.0 chemiluminescent substrate (Cyanagen) or SuperSignal West Femto Maximum Sensitivity Substrate (34095; Thermo Fisher) at ChemiDoc MP Imaging system (Bio-Rad).

**Table 4:**
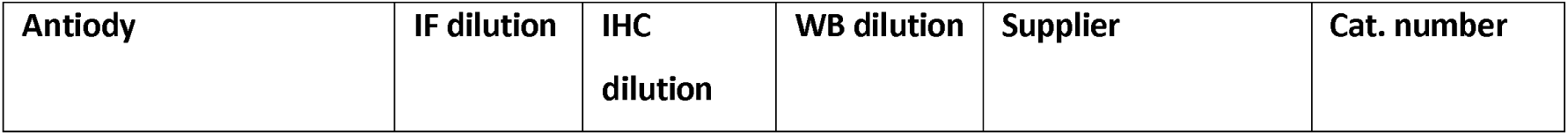
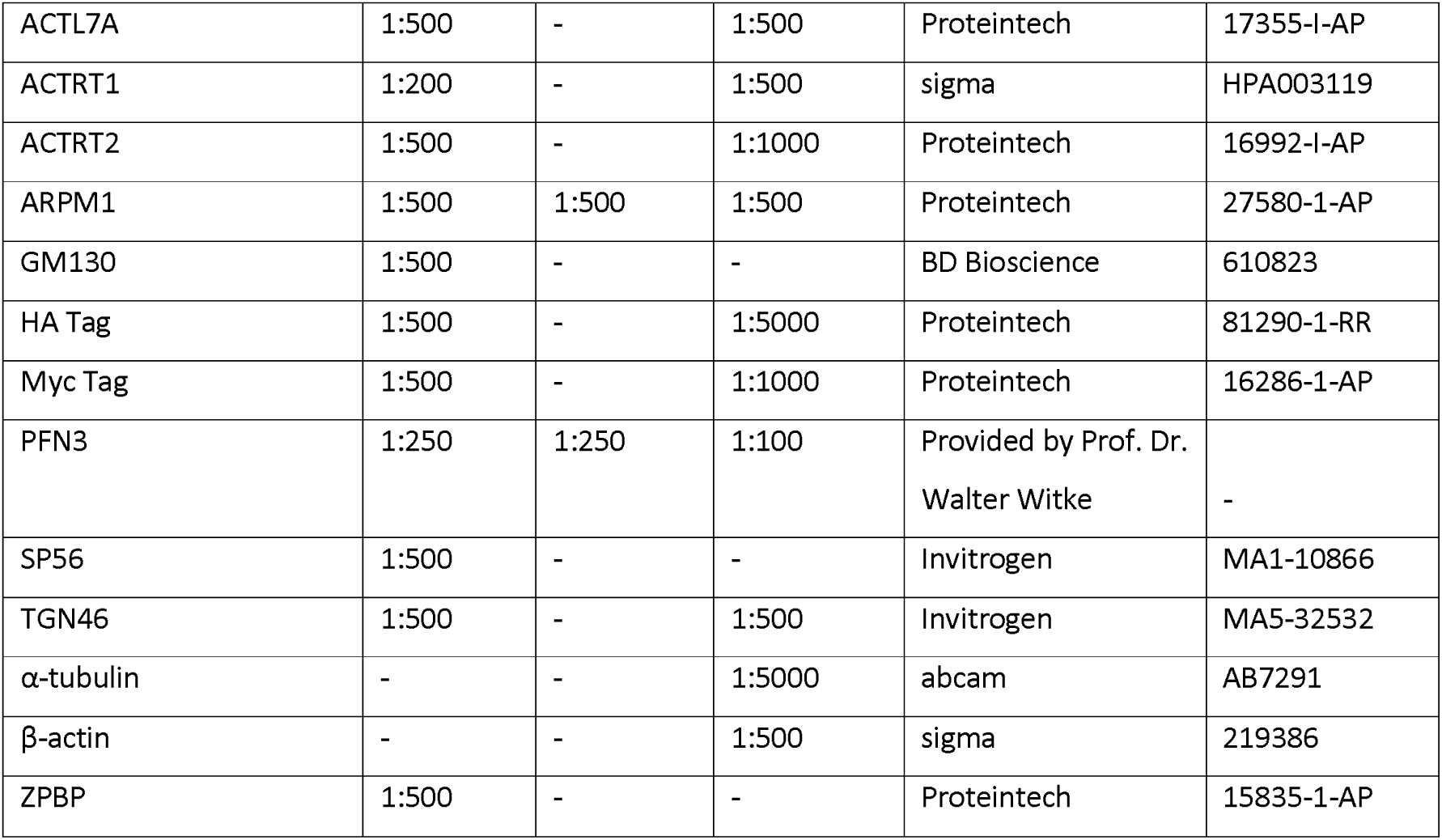
Antibody dilutions.

### Immunofluorescent stainings

For IF stainings, Bouin-fixed testis tissue sections were deparaffinized in xylene and rehydrated in decreasing alcohol, while mature sperm cells isolated from cauda epididymis were fixed with methanol and acetic acid (3:1), dropped on glass slides, and air dried. For tissue sections, heat-activated antigen retrieval was performed using citrate buffer (pH 6.0). All samples were first washed in PBS twice, and permeabilized using 0.1% Triton X-100 for 10 min at room temperature. Next, the samples were blocked with 5% BSA for 30 min, followed blocking with normal horse serum (Vectorlabs, Burlingame, CA, USA; DI-1788) for 30 min at room temperature. All primary antibodies were incubated overnight at 4°C at dilutions listed in Table 4. The secondary antibodies were incubated for 1 h at room temperature using VectaFluor Labeling Kit DyLight 488 or DyLight 594 (Vectorlabs, Burlingame, CA, USA; DI-1788, DI-1794). After washing with PBS two times, slides were mounted with DAPI containing mounting medium (ROTImount FluorCare DAPI, Carl Roth; HP20.1).

For the analysis of acrosome morphology, PNA-FITC Alexa Fluor 488 conjugate (Molecular Probes, Invitrogen, Waltham, MA, USA; L21409) was used on the Bouin fixed testis tissues and epididymal sperm fixed in paraformaldehyde 4%, for 20 min at room temperature. Permeabilization and blocking were performed as described above and the samples were incubated with PNA-FITC 5 μg/ml for 30 min at room temperature. Finally, the slides were mounted using DAPI containing mounting medium. For the assessment of chromatin compaction by chromomycin A3 (CMA3), 10 µL of epididymal sperm fixed in methanol and acetic acid (3:1) was dropped on glass slides and air dried. The slides were covered in CMA3 solution and incubated for 20 minutes in a humid chamber at room temperature.

Slides were then rinsed in PBS twice and mounted with DAPI containing mounting medium (ROTImount FluorCare DAPI, Carl Roth; HP20.1). All stainings were performed on a minimum of three animals per genotype.

For immunofluorescence staining HEK293T cells were grown on glass coverslips in a 24 well plate. Cells were fixed in 4% PFA for 15 minutes, 24 h after transfection and washed in PBS. After permeabilization with 0.3% Triton X-100 for 10 minutes, cells were washed in PBS and blocked with Normal Horse Serum for 20 minutes, followed by blocking with 5% BSA for 30 minutes. Primary antibodies were incubated at 4°C overnight at dilutions listed in Table 4. Next, after washing in PBS, cells were incubated with secondary horse anti-rabbit antibody DyLight™ 594 (DI-1094, Vectorlabs, Burlingame, CA, USA) for 1 h at room temperature. To counterstain the nuclei, cells were incubated with 0.01 mg/ml Hoechst (bisBenzimide H 33342, Sigma-Aldrich) for 10 minutes. Finally, after PBS wash, coverslips were mounted on glass slides using Fluoroshield™ (F6182, Sigma-Aldrich).

### Transmission Electron Microscopy

For TEM analysis sperm cells and testis samples from all three genotypes were fixed with 4% PFA and 2.5% glutaraldehyde in PBS overnight. After washing in cacodylate buffer, the samples were incubated with 1% osmium tetroxide and 0.8% ferricyanate in cacodylate buffer for 2 h. Samples were washed again with cacodylate buffer and pelleted at 600 g before dehydration in ethanol row (30%, 50%, 70%, 90%, 95%, 100%), and propylene oxide. The dehydration step with 70% ethanol was done in the presence of 0.5% uranyl acetat for 1 h. Samples were then infiltrated with propylene oxide/Epon mixtures (1:1, 1:2) before infiltration with pure Epon. Epon blocks were incubated at 60°C for 48 h. 70 nm ultrathin sections were cut with an ultramicrotome (RMC Boeckeler Powertome) and collected on formvar-carbon-coated TEM copper slot girds. TEM grids were counterstained with uranyl acetate and lead citrate before imaging transmitted electrons with a Crossbeam 550 (Zeiss) and a retractable STEM detector. SmartSEM software (Zeiss) was used to acquire images at 30 kV acceleration voltage with 150 pA current in imaging mode.

### Plasmids

Plasmids used in this study were generated by amplifying *Arpm1*, *Actrt1*, *Actrt2*, *Actl7a*, *Pfn3*, *Spem2* and *ZPBP* from C57Bl/6J mouse testis cDNA using overhang primers which introduced suitable restriction enzyme motifs (Supplementary Table S2). *Arpm1* was cloned in pCMV_HA-C Clonetech (635690) vector, while *Actrt1*, *Actrt2*, *Actl7a*, *Pfn3*, *Spem2* and *Zpbp* were cloned in pCMV_Myc-C Clonetech (635689). Correct sequence and insertion were verified by sequencing.

### Transfection

HEK293T (RRID: CVCL_0063) cells were cultured in DMEM (Gibco) with 10% FBS supplemented with antibiotics and transfected at 60-70% confluency. Transfection/Co-transfection was performed using 2.5 μg of expression plasmid with Lipofectamin™ 3000 according to manufacturer’s instructions. As control for transfection efficiency pEGFP_N3 plasmid was transfected. After 48 h since transfection, cells were imaged using a Leica DM5500 B microscope and proteins were isolated for co-IP analysis. HEK293T cells were regularly checked for the presence of mycoplasma contamination.

### Co-Immunoprecipitation

For Co-immunoprecipitation either Pierce® HA IP Kit or Pierce® c-Myc Tag IP Kit was used according to manufacturer’s instructions. Briefly, 200 μl lysate with anti-HA agarose was added to a provided spin column and incubated ON at 4°C. Next, spin columns were placed in a collection tube and the flow through was ensured through 10 s pulse centrifugation. After three washing steps in TBS-T (0.05% Tween 20), elution was performed by adding 25 μl 2x non-reducing sample Buffer, incubation at 95°C for 5 min and followed by pulse centrifugation for 10 s. For further analysis input, flow through and eluate were used and analyzed by western blotting.

### Evolutionary analysis

The evolutionary rate of *ARPM1* among rodents and primates was analyzed according to Lüke et al. (2016). *ARPM1* coding sequences for 33 rodent and 35 primate species, as well as *Bos taurus* and *Canis lupus familiaris* as outgroups, were obtained from NCBI genbank and Ensembl genome browser. Phylogenetic trees of the included species were built according to the “Tree of Life web project” (Letunic and Bork, 2021) The webPRANK software was applied for codon-based phylogeny-aware alignment of orthologous gene sequences. The tree and alignment were visualized using the ETE toolkit (Löytynoja and Goldman, 2010). To evolutionary rates and selective pressures were determined using codeML implemented in PAML4.9 (Yang, 2007). The evolutionary rate is based on the calculation of the nonsynonymous/synonymous substitution rate ratio (ω = dN/dS). It distinguishes between purifying selection (ω<1), neutral evolution (ω=1) and positive selection (ω>1). Different null and alternative models (M) were applied. The M0 model served as the basic model for all performed analyses. Different codon frequency settings were tested for the M0 model of each gene and the setting with the highest log likelihood was chosen. To test whether alternative models describe the selective constraints within a dataset better than the null models, likelihood-ratio-tests (LRT) were performed. In order to determine the overall evolutionary rate and selective pressure on the coding sequence among all included species we employed two models: M0 “one ratio” in which all branches were constrained to evolve at the same freely estimated evolutionary rate; M0fix (fixed ratio) in which the evolutionary rate for all branches was constrained to ω=1. The M0 model calculates the overall evolutionary rate. An LRT between M0 and M0fix was performed to determine the calculated evolutionary rate. In order to determine if the selective pressures differ between rodents and primates, we employed two models: M0 “one ratio” and MC “two-ratio” which allows for the estimation of a free and independent ω for the marked clades. To test if the alternative MC model presents a better fit for the data, we compared the models log likelihood values by LRT. To test the evolutionary rate along coding sequences and infer codon sites under positive or purifying selection we applied LRT comparing the null model M1a “nearly neutral”, which does not allow sites with ω>1 with the alternative model M2a “selection” which does. The models assign the codon sites into different classes: Class 0: sites under purifying selection (0>ω>1); Class 1: sites evolving neutrally (relaxed constraint) (ω = 1); Class 2a (only M2a): sites subject to positive selection (ω>1). Bayesian statistics were used to identify those codons that have been subject to either positive selection or purifying selection. Only sites with posterior probabilities (Naïve Empirical Bayes and Bayes Empirical Bayes) higher than 0.95 to be assigned to class 0 or class 2a were determined to be under purifying or positive selection respectively.

### Statistics

For all analyses, statistical significance was calculated using a two-tailed, unpaired Student’s *t*-test. *P*<0.05 was considered significant (\**P*<0.05, \*\**P*<0.005, \*\*\**P*<0.001, \*\*\*\**P*<0.0001). If not indicated otherwise, values were given as mean values with standard deviation.

## Acknowledgment

This study was supported by a grant from the German Research Foundation (DFG Scho503/29-1, project # 546839078) to HS. We are grateful to Greta Zech, Angela Egert, Andrea Jäger and Gaby Beine for excellent technical assistance. We would like to thank the Core Facility for Microscopy of the Medical Faculty at the University of Bonn for providing support and instrumentation funded by the Deutsche Forschungsgemeinschaft (DFG, German Research Foundation, project number: 388171357).

## Author Contributions

A.K, G.E.M., N.U. and H.S. conceived and designed the experiments. N.U. generated the *Arpm1*-deficient mouse model. A.K, S.S, E.O. L.D.H and G.E.M. conducted the experiments. A.K. and H.S. were major contributors in writing the manuscript. All authors read and approved the final manuscript.

## Data availability

All relevant data and resources can be found within the article and its supplementary information.

