## Supplementary material for "Actin-related protein M1 (ARPM1) required for acrosome biogenesis and sperm function in mice": Fig. S1

### Figure S1

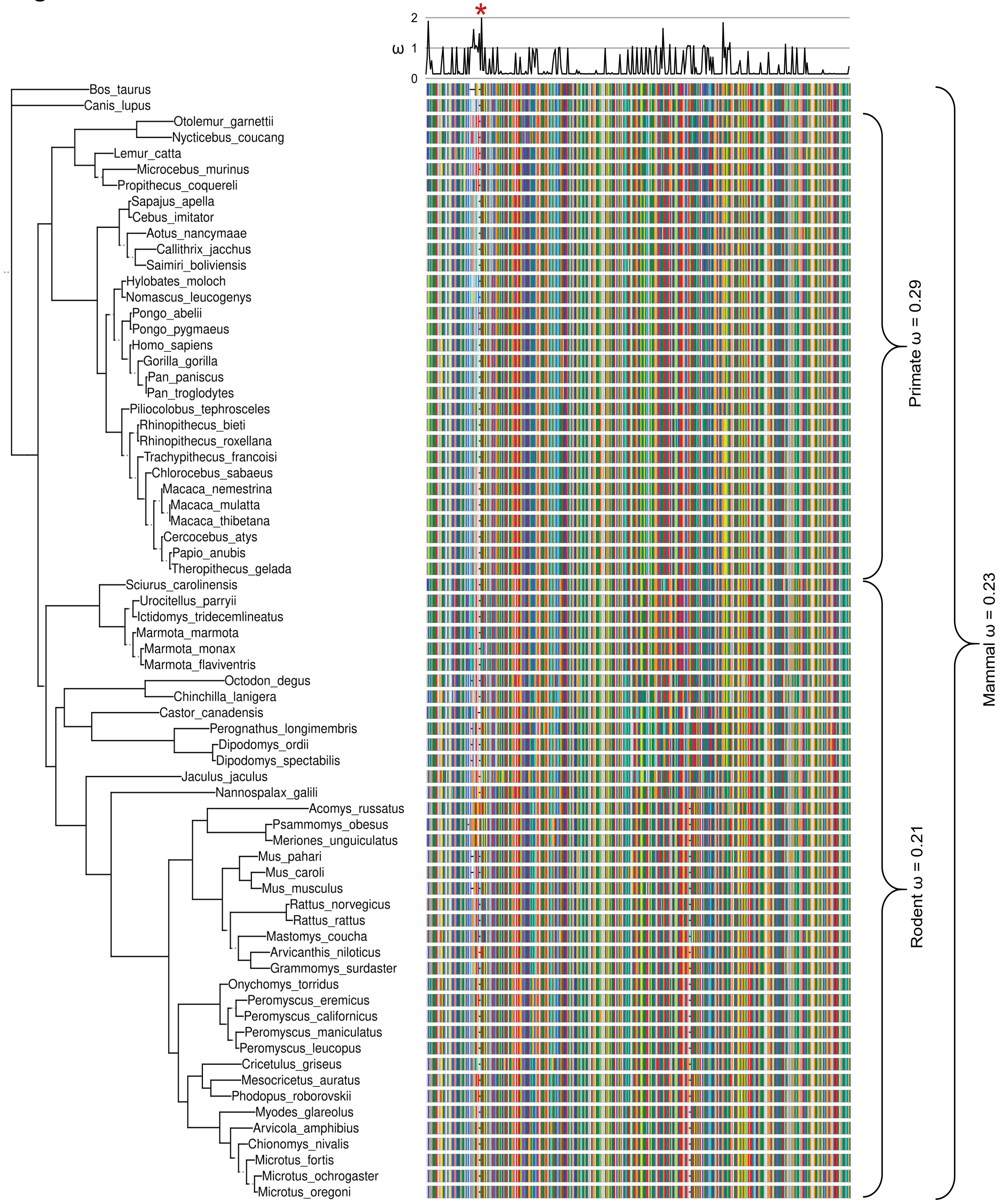


**Figure S1: Arpm1 is evolutionarily conserved**
Phylogeny of included species, with branch length denoting the number of nucleotide substitutions per codon, and a schematic depiction of the ARPM1 amino acid alignment utilized in the PAML CodeML analysis. Evolutionary rate (ω) obtained by CodeML model M0 is shown for the whole alignment (Mammal ω) and the primate (Primate ω) and rodent clade (Rodent ω). The graph at the top depicts ω per codon site across the whole tree (CodeML model M2a). Asterisks indicate significantly positively selected sites. See also: Table S1.
