## Supplementary material for "Actin-related protein M1 (ARPM1) required for acrosome biogenesis and sperm function in mice": Fig. S2

### Figure S2

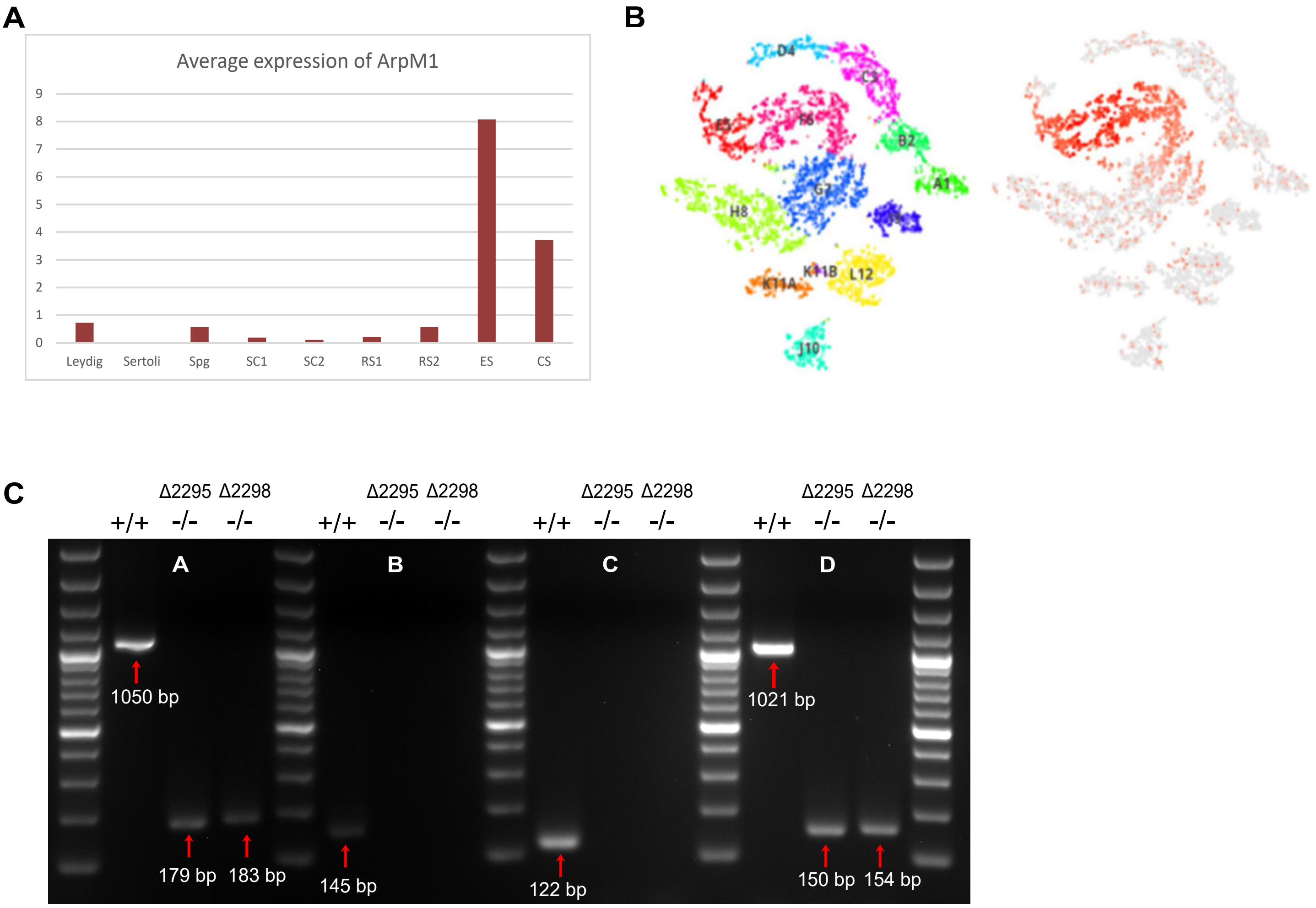


**Figure S2:**
(A) Average expression of Arpm1 in cell populations within adult mouse testis: Spg – spermatogonia; SC1 – spermatocytes type 1; SC2 – spermatocytes type 2; RS1 – early round spermatids; RS2 – late round spermatids; ES – elongating spermatids; CS – compacted sperm.
(B) Average expression of Arpm1 (right panel) in human testis populations extracted from Human Testis Atlas. Clusters of cells are shown in the left panel: A1 – spermatogonial stem cells; B2 – spermatogonial stem cells; C3 – early spermatids; D4 – late spermatids; E5 – round spermatids; F6 - elongated spermatids; G7 – spermatocytes; H8 – spermatocytes; I9 – macrophages; J10 – Endothelia; K11A – Myoid; L12 – Leydig cells
(C) PCR on testicular cDNA from WT and Arpm1-/- mice (Δ2295 and Δ2298). Band sizes are shown for each PCR product.
