## Supplementary material for "Actin-related protein M1 (ARPM1) required for acrosome biogenesis and sperm function in mice": Fig. S3

### Figure S3

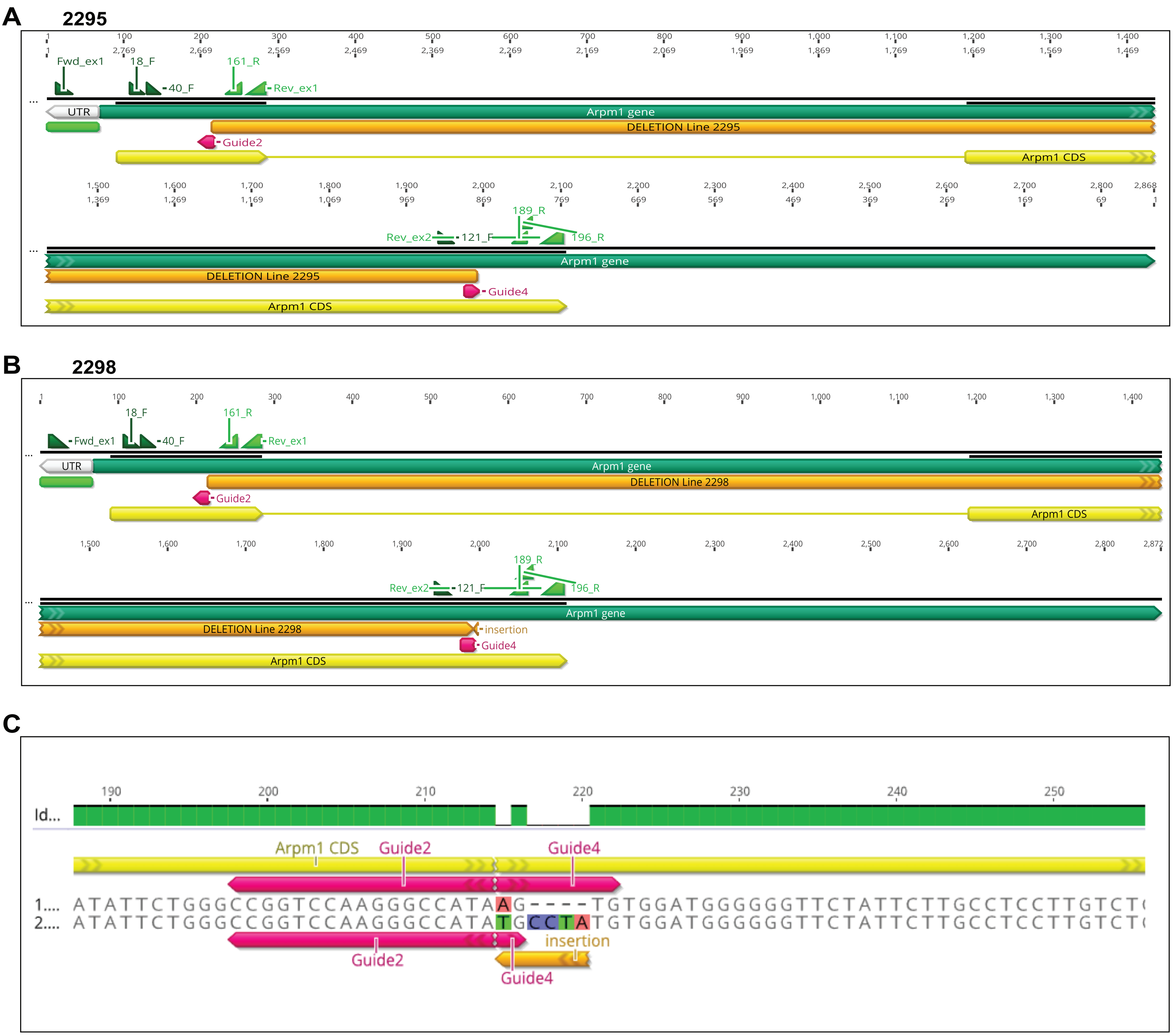


**Figure S3:**
A) Schematic representation of *Arpm1* gene sequence from Δ2295 line (yellow) compared to WT Arpm1 gene sequence (green). Deleted region is shown in orange. Primers used for genotyping PCR and PCR on cDNA (green), as well as guide sequences (red) are shown in respective positions.
B) Schematic representation of *Arpm1* gene sequence from Δ2299 line (yellow) compared to WT Arpm1 gene sequence (green). Deleted region is shown in orange. Primers used for genotyping PCR and PCR on cDNA (green), as well as guide sequences (red) are shown in respective positions.
C) Amino acid sequence differences between Δ2295 line (upper row) and Δ2298 line (lower row). Guide sequences are shown in magenta, while orange depicts an insertion detected in Δ2298 line.
