## Supplementary material for "Actin-related protein M1 (ARPM1) required for acrosome biogenesis and sperm function in mice": Fig. S4

### Figure S4

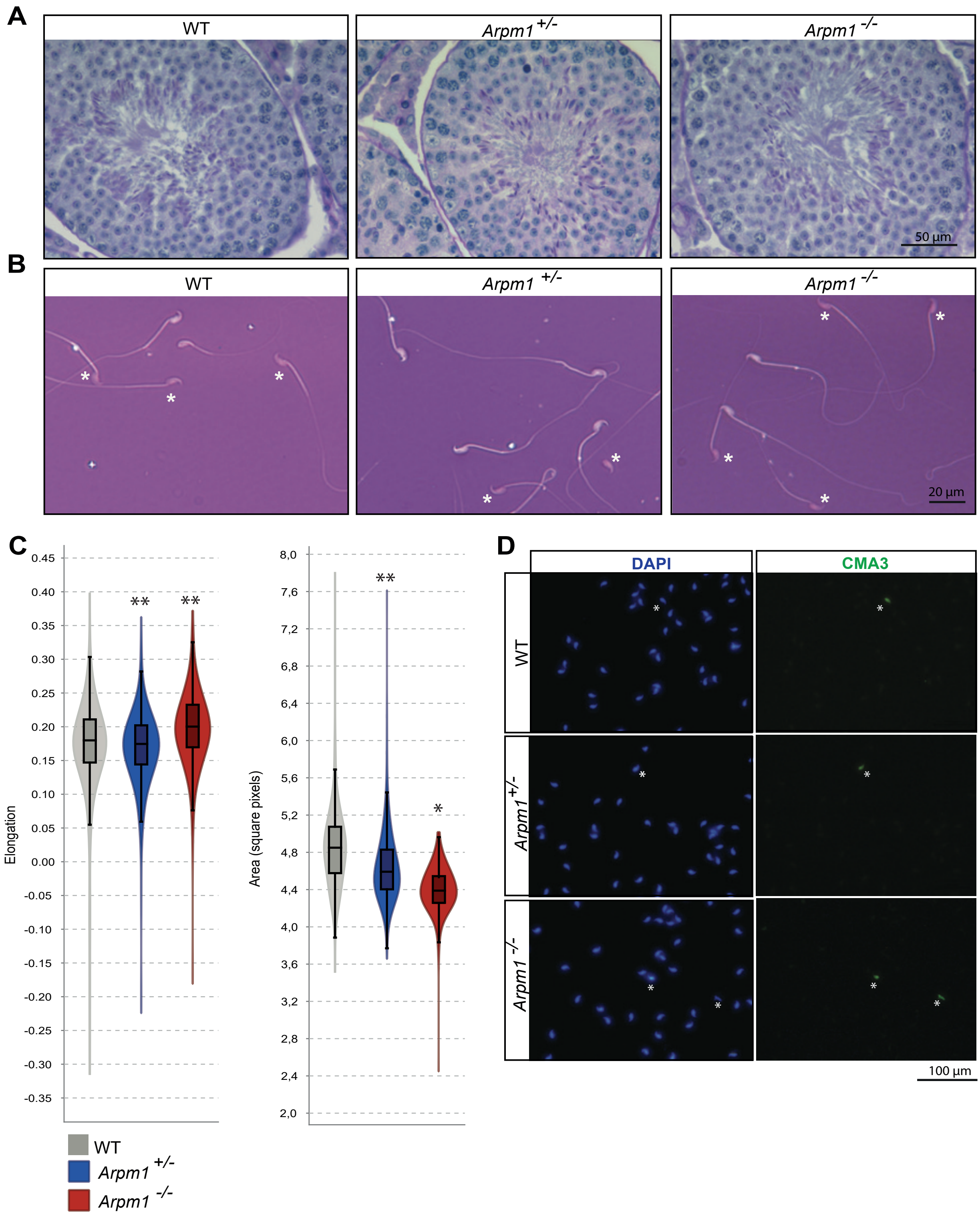


**Figure S4:**
(A) PAS staining of testicular tissue sections from WT, *Arpm1+/-* and *Arpm1-/-* mice depicting all cell populations of the testis. Scale bar: 50 µm.
(B) Representative images of Eosin-Nigrosin staining of epididymal sperm from WT, *Arpm1+/-* and *Arpm1-/-* mice. Stained cells depicted with white asterisks were counted as EN positive while white cells were counted as EN negative. Scale bar: 20 µm.
(C) Elongation and area of nuclei from WT, *Arpm1+/-* and *Arpm1-/-* sperm. The minimum detection area was set to 1000 pixels, while the maximum detection area was 7000 pixels. (*P<0.05, **P<0.005, ***P<0.001, ****P<0.0001)
(D) Epididymal sperm from WT, *Arpm1+/-* and *Arpm1-/-* mice stained with CMA3 (right panel) and counterstained with DAPI (left panel). CMA3 positive nuclei are depicted with white asterisks. Scale bar: 100 µm.
