## Supplementary material for "Actin-related protein M1 (ARPM1) required for acrosome biogenesis and sperm function in mice": Fig. S5

### Figure S5

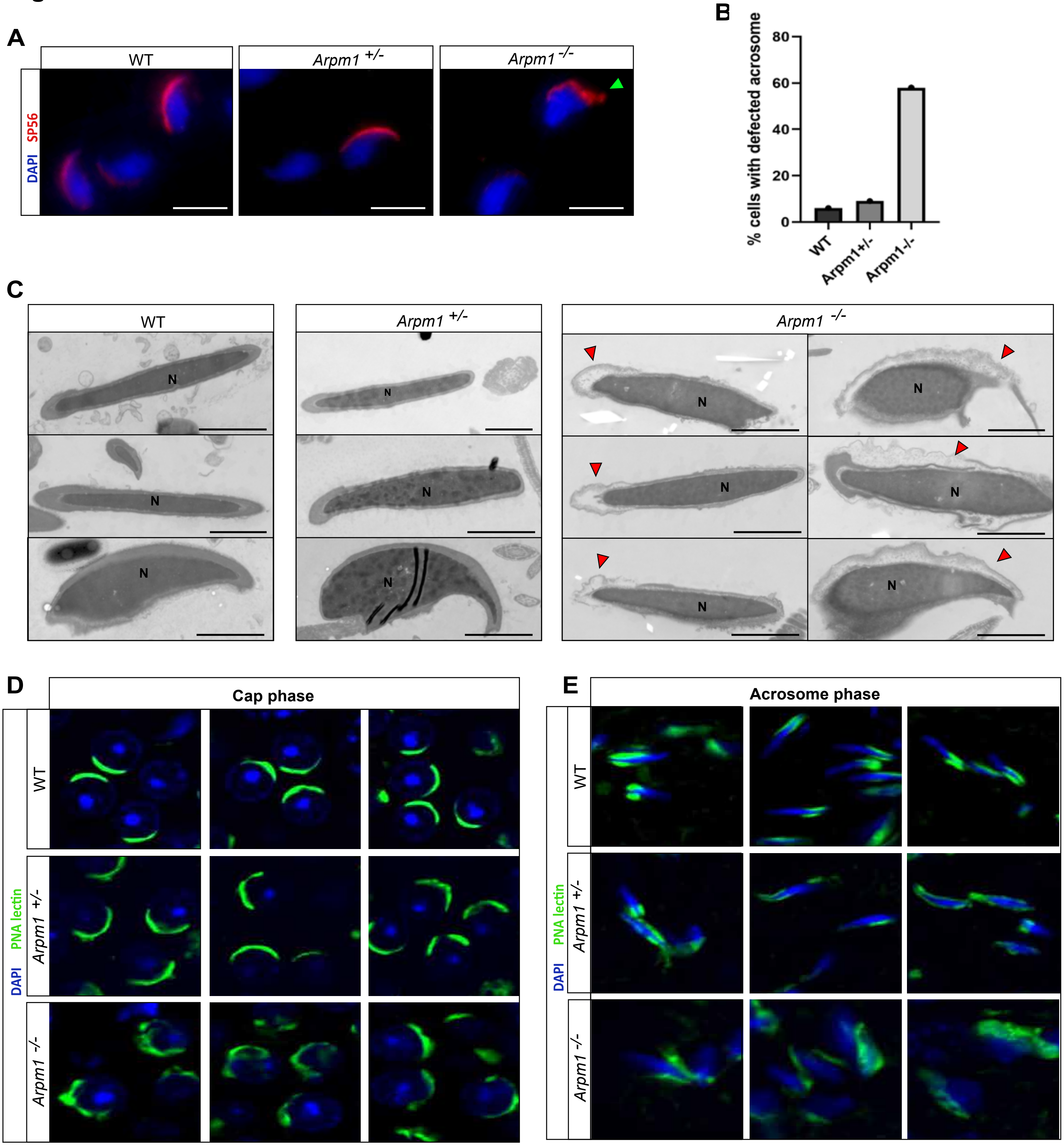


**Figure S5:**
(A) Immunofluorescence staining against acrosome marker SP56 in epididymal sperm of WT, *Arpm1+/-* and *Arpm1-/-* male mice (red). The green arrowhead depicts the damaged acrosome. Nuclei were counterstained with DAPI. Staining was performed on three animals from each genotype. Scale bar: 10 µm.
(B) Quantification of acrosomal defects observed in SP56 stained epididymal sperm from WT, *Arpm1+/-* and *Arpm1-/-* male mice. At least 100 cells were counted per genotype.
(C) Transmission electron microscopy (TEM) micrographs of WT, *Arpm1+/-* and *Arpm1-/-* epididymal sperm. Defected acrosomes in Arpm1-/- sperm are depicted with red arrowheads. N: nucleus. Scale bar: 5 µm.
(D) PNA-lectin immunofluorescence staining of the acrosome in testicular tissue sections of WT, *Arpm1+/-* and *Arpm1-/-* male mice (green). Single round spermatids in cap phase are depicted. Nuclei were counterstained with DAPI. Staining was performed on three animals from each genotype. Scale bar: 5 µm.
(E) PNA-lectin immunofluorescence staining of the acrosome in testicular tissue sections of WT, *Arpm1+/-* and *Arpm1-/-* male mice (green). Elongating spermatids in acrosome phase are depicted. Nuclei were counterstained with DAPI. Staining was performed on three animals from each genotype. Scale bar: 5 µm.
