## Supplementary material for "Actin-related protein M1 (ARPM1) required for acrosome biogenesis and sperm function in mice": Fig. S6

### Figure S6

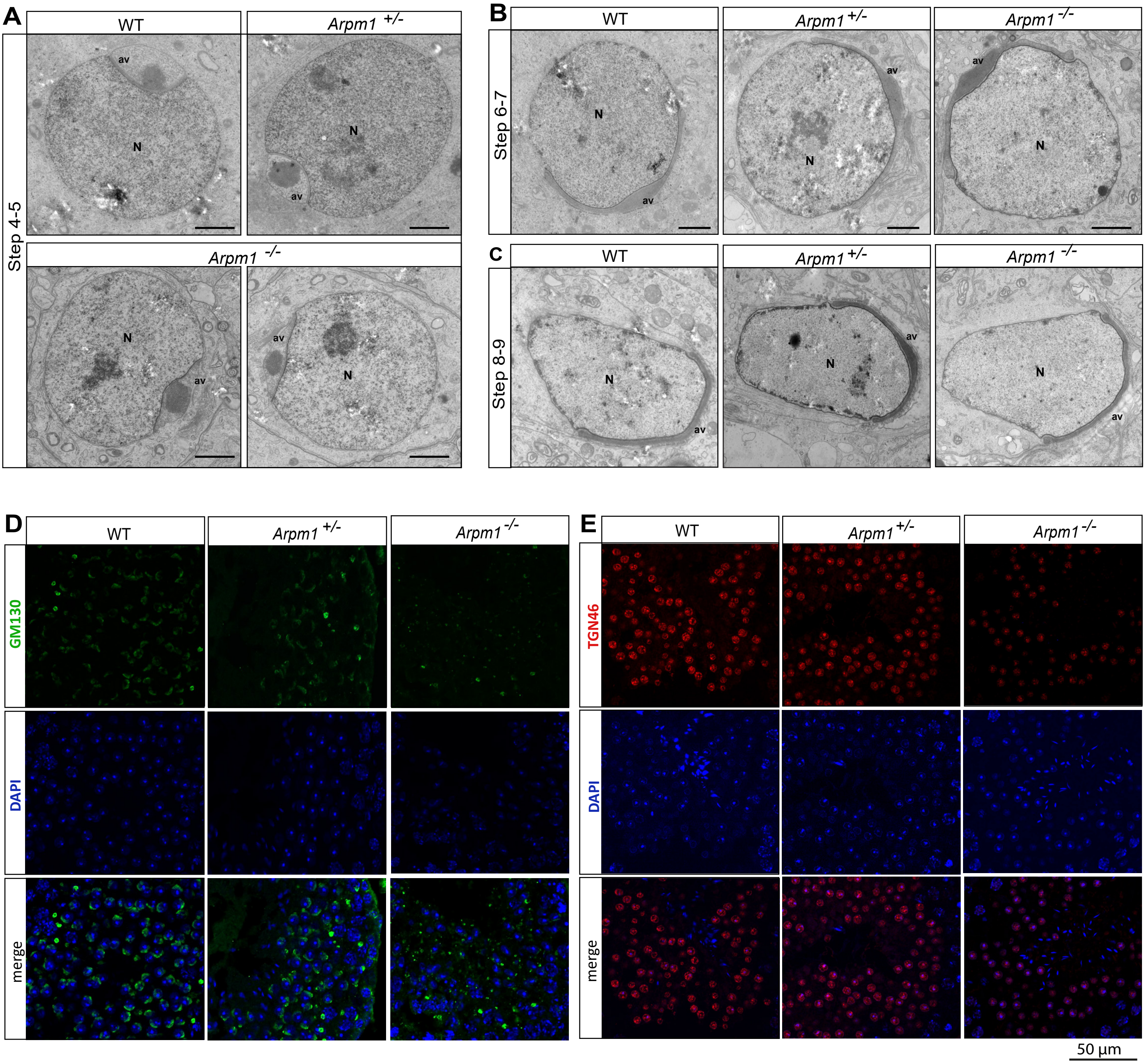


**Figure S6:**

(A) TEM micrographs of WT, *Arpm1+/-* and *Arpm1-/-* round spermatids in step 4-5 at Golgi phase of acrosome development. N: nucleus; av: acrosomal vesicle. Scale bar: 1 µm

(B) TEM micrographs of WT, *Arpm1+/-* and *Arpm1-/-* round spermatids in Cap phase of acrosome development. N: nucleus; av: acrosomal vesicle. Scale bar: 1 µm

(C) TEM micrographs of WT, *Arpm1+/-* and *Arpm1-/-* elongating spermatids in acrosome phase of acrosome development. N: nucleus; av: acrosomal vesicle. Scale bar: 1 µm

(D) Immunofluorescence staining of testicular tissue sections from WT, *Arpm1+/-* and *Arpm1-/-* male mice against GM130 (green). In WT and *Arpm1+/-* round spermatids, GM130 signal forms a cap-like structure along the developing acrosome. In *Arpm1-/-* round spermatids GM130 signal was scattered irregularly. Nuclei were counterstained with DAPI (blue). Scale bar: 50 µm

(E) Immunofluorescence staining of testicular tissue sections from WT, *Arpm1+/-* and *Arpm1-/-* male mice against TGN46 (red). In *Arpm1-/-* round spermatids TGN46 signal was less intense compared to WT and *Arpm1+/-* round spermatids. Nuclei were counterstained with DAPI (blue). Scale bar: 50 µm
