## Supplementary material for "Actin-related protein M1 (ARPM1) required for acrosome biogenesis and sperm function in mice": Fig. S7

### Figure S7

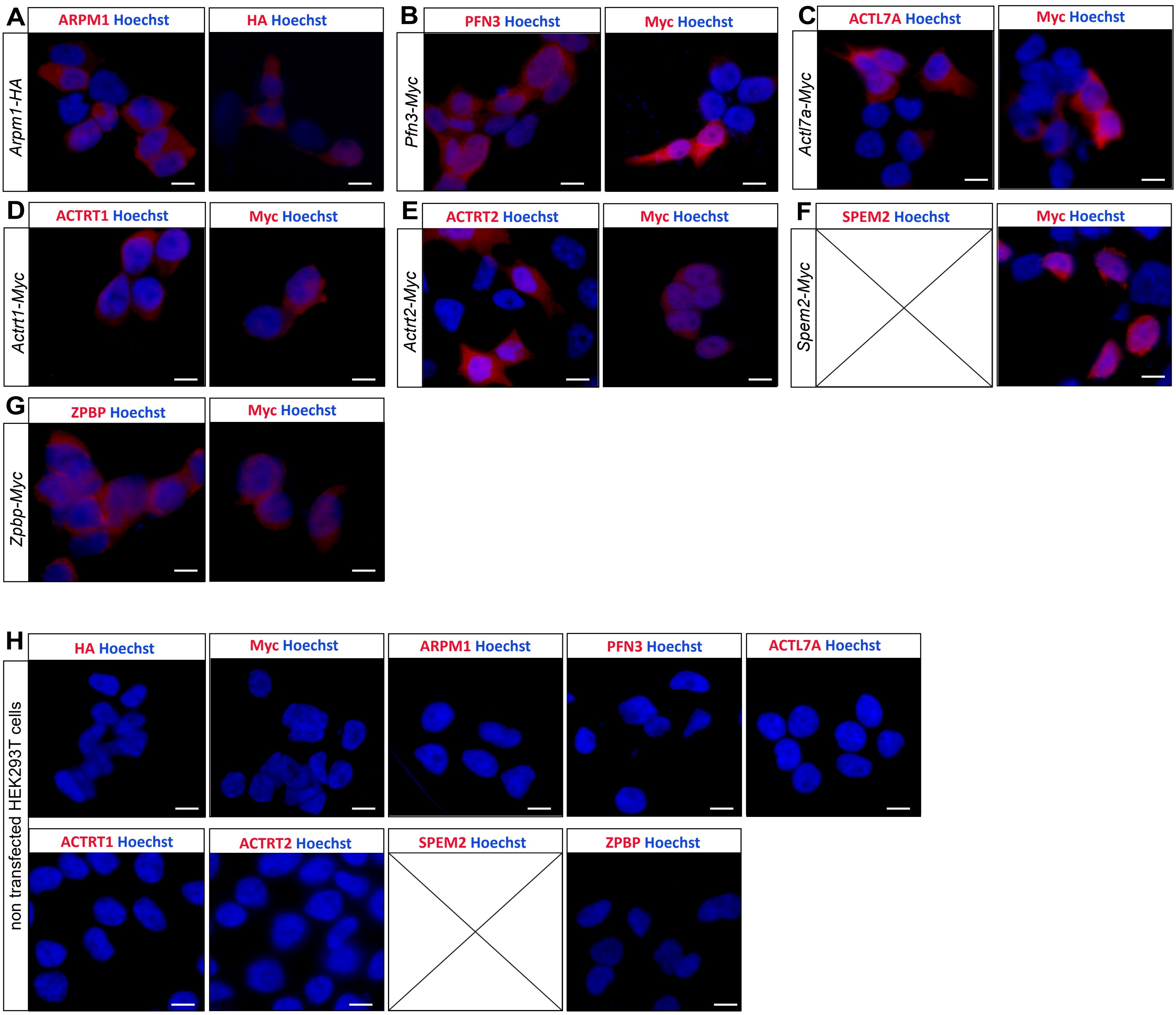


**Figure S7:**
(A-G) Immunofluorescent staining of transfected HEK293T cells: (A) *Arpm1*-HA transfected HEK293T cells stained against ARPM1 (left panel, red) and HA (right panel, red); (B) *Pfn3*-Myc transfected HEK293T cells stained against PFN3 (left panel, red) and Myc (right panel, red). The signal is detectable in the cytoplasm as well as in the nucleus; (C) *Actl7a*-Myc transfected HEK293T cells stained against ACTL7A (left panel, red) and Myc (right panel, red); (D) *Actrt1*-Myc transfected HEK293T cells stained against ACTRT1 (left panel, red) and Myc (right panel, red); (E) *Actrt2*-Myc transfected HEK293T cells stained against ACTRT2 (left panel, red) and Myc (right panel, red); (F) *Spem2*-Myc transfected HEK293T cells stained against Myc (right panel, red); (G) *Zpbp*-Myc transfected HEK293T cells stained against ZPBP (left panel, red) and Myc (right panel, red); Nuclei were counterstained with Hoechst 3458. Scale bar: 10 µm
(H) Immunofluorescent staining against HA-tag, Myc-tag, ARPM1, PFN3, ACTL7A, ACTRT1, ACTRT2 and ZPBP on non-transfected HEK293T cells. Nuclei were counterstained with Hoechst 3458. Scale bar: 10 µm
