## Supplementary material for "Actin-related protein M1 (ARPM1) required for acrosome biogenesis and sperm function in mice": Fig. S8

### Figure S8

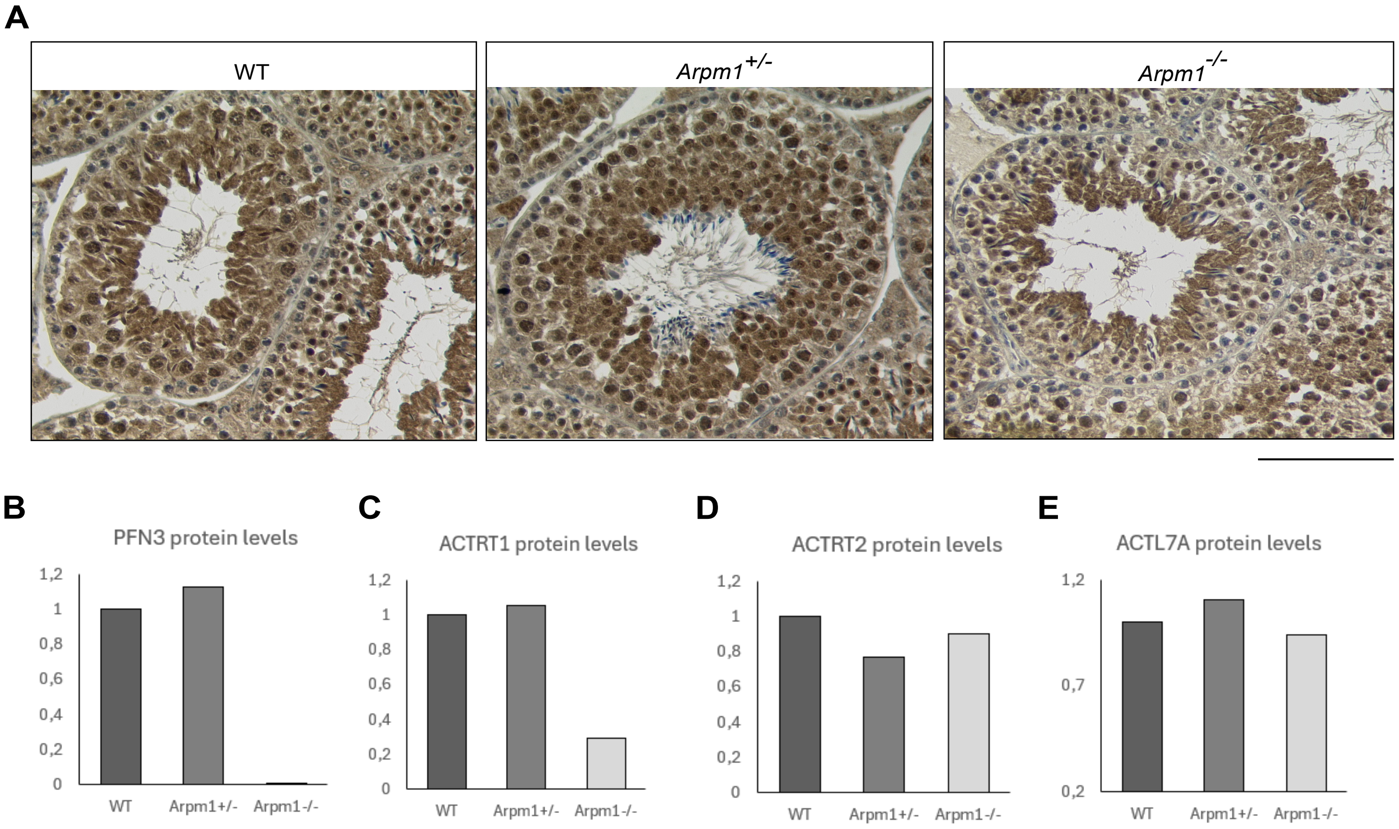


**Figure S8:**
A) Immunohistochemical staining of testicular tissue sections from WT, *Arpm1+/-* and *Arpm1-/-* mice against PFN3. Scale bar: 100 µm
B-E) Quantification of protein levels of B) PFN3; C) ACTRT1 D) ACTRT2 E) ACTL7A determined by immunoblotting and normalized to α-tubulin.
