## Supplementary Table S 1 for "Actin-related protein M1 (ARPM1) required for acrosome biogenesis and sperm function in mice"

| **Selection on:** | **Gene / Clade** | **LnL null model** | | **LnL alternative model** | **LRT** | **p** | **ω** | **interpretation** |  |  |  |  |
| --- | --- | --- | --- | --- | --- | --- | --- | --- | --- | --- | --- | --- |
| whole sequence, whole tree | *Arpm1 (Actrt3)* | 13626.525 (M0) | | 14179.431 (M0fix) | -1105,81 | n.s. | 0.23 (M0) | overall conserved |  |  |  |  |
| whole sequence, clades / lineages | *Arpm1 (Actrt3)* | Primates | 13626.525 (M0) | 13619.801 (MC) | 13,45 | 0,01 | 0.29 (MC) | both lineages conserved |  |  |  |  |
|  |  | Rodentia |  |  |  |  | 0.21 (MC) |  |  |  |  |  |
| codon sites, whole tree | *Arpm1 (Actrt3)* | 13306.638 (M1a) | | 13302.419 (M2a) | 8,43 | 0,05 | n.a. | **prop 0** | **prop 1** | **prop 2a** | **PSS** | **PUR** |
|  |  |  |  |  |  |  |  | 0,75 | 0,24 | 0,01 | 50Q | 37R,62R,68S,73R,  103V,106T,116R,  129V,175V,288R,  302T,310R,356T,  369H,371R,  148T,206R,276C,  294N,366N,367I,  368V,372C,11D,  23G,27P,33N,  81D,158G,164P,  242D,243G,273D,  284D,322N,327V,  330P,352D,363V,  25R,1M,83E,88H,  214E,218Y,257E,  299G,333R,342S,  348S,353M |

**Table S1:** **Results of the evolutionary analysis using CodeML (PAML4.9)** LnL = log Likelihood value of the model; LRT = Likelihood Ratio Test; p = p-value of the LRT; ω = evolutionary rate (dN/dS); prop = proportion of sites assigned to the respective site class; PSS = positively selected sites; PUR = sites under purifying selection / conserved sites; for explanation of the models see M&M.
