## Supplementary Table S2 for "Actin-related protein M1 (ARPM1) required for acrosome biogenesis and sperm function in mice"

**Table S2: Overhang primers and plasmids**

| **Primer name** | **Sequence (5´-3´)** | |
| --- | --- | --- |
| OV fwd_EcoR1_ACTRT1 | AAAAGAATTCATGCTTGATCCAGCTAGATTAGAT | |
| OV rev_XhoI_ACTRT1 | TTTTCTCGAGAAAAGCATTTTCTTTGAACCACAAATG | |
| OV fwd_EcoR1_ACTRT2 | AAAAGAATTCATGTTTAACCCACTGGTACTAGATTC | |
| OV rev_XhoI_ACTRT2 | TTTTCTCGAGAGAAGCACCGCCTCTGGAC | |
| OV_fwd_EcoRI_ACTL7A | AAAAGAATTCATGTCTCTGGATGGTGTGTG | |
| OV_rev_KpnI_ACTL7A | TTTTGGTACCGAAGCACCTTCTGTAGAGGA | |
| OV_fwd_ApaI_ARPM1 | AAAAGGGCCCATGAGCGGCTACCAGCCAC | |
| OV_rev_KpnI_ARPM1 | TTTTGGTACCAAAGCATCTCTGGTGTACAATGTTGG | |
| OV_fwd_EcoRI_PFN3 | AAAAGAATTCATGAGTGACTGGAAGGGCTACATCAGTGC | |
| OV_rev_KpnI_PFN3 | TTTTGGTACCAGAGCACTGCTCACGCAGCCCACCAA | |
| OV_fwd_EcoRI_ZPBP | AAAAGAATTCATGTGTACCCAAGCTGGGTCCCAA | |
| OV_fwd_EcoRI_SPEM2 | AAAAGAATTCATGGAAAACCAGCTGTGGCAGAACA | |
| OV_rev_XhoI_SPEM2 | TTTTCTCGAGAGTTAAGTTTCCCTGTCCGGCTTCTA | |
| **Plasmid name** | | **Source** |
| pEGFP_N3 | | Clonetech (6080-1) |
| pCMV_HA-C | | Clonetech (635690) |
| pCMV_Myc-C | | Clonetech (635689) |
| pCMV_Myc-C_ACTRT1 | | Generated in this study |
| pCMV_Myc-C_ACTRT2 | |  |
| pCMV_Myc-C_ACTL7A | |  |
| pCMV_Myc-C_PFN3 | |  |
| pCMV_Myc-C_ZPBP | |  |
| pCMV_Myc-C_SPEM2 | |  |
